# *In vivo* human neurite exchange imaging (NEXI) at 500 mT/m diffusion gradients

**DOI:** 10.1101/2024.12.13.628450

**Authors:** Kwok-Shing Chan, Yixin Ma, Hansol Lee, José P. Marques, Jonas Olesen, Santiago Coelho, Dmitry S Novikov, Sune Jespersen, Susie Y. Huang, Hong-Hsi Lee

**Author notes:** Correspondence to: Hong-Hsi Lee, MD, PhD, Athinoula A. Martinos Center for Biomedical Imaging, Charlestown, MA, United States, 149 13^th^ St, Charlestown, MA, USA.

## Abstract

Evaluating tissue microstructure and membrane integrity in the living human brain through diffusion-water exchange imaging is challenging due to requirements for a high signal-to-noise ratio and short diffusion times dictated by relatively fast exchange processes. The goal of this work was to demonstrate the feasibility of *in vivo* imaging of tissue micro-geometries and water exchange within the brain gray matter using the state-of-the-art Connectome 2.0 scanner equipped with an ultra-high-performance gradient system (maximum gradient strength=500 mT/m, maximum slew rate=600 T/m/s). We performed diffusion MRI measurements in 15 healthy volunteers at multiple diffusion times (13-30 ms) and *b*-values up to 17.5 ms/μm^2^. The anisotropic Kärger model was applied to estimate the exchange time between intra-neurite and extracellular water in gray matter. The estimated exchange time across the cortical ribbon was around (median±interquartile range) 13±8 ms on Connectome 2.0, substantially faster than that measured using an imaging protocol compatible with Connectome 1.0-alike systems on the same cohort. Our investigation suggested that the NEXI exchange time estimation using a Connectome 1.0 compatible protocol was more prone to residual noise floor biases due to the small time-dependent signal contrasts across diffusion times when the exchange is fast (≤20 ms). Furthermore, spatial variation of exchange time was observed across the cortex, where the motor cortex, somatosensory cortex and visual cortex exhibit longer exchange times compared to other cortical regions. Non-linear fitting for the anisotropic Kärger model was accelerated 100 times using a GPU-based pipeline compared to the conventional CPU-based approach. This study highlighted the importance of the chosen diffusion times and measures to address Rician noise in dMRI data, which can have a substantial impact on the estimated NEXI exchange time and require extra attention when comparing NEXI results between various hardware setups.

**Highlights:**

- The Connectome 2.0 scanner equipped with 500 mT/m gradients enables high-sensitivity diffusion MRI for imaging exchange times *in vivo*
- The global exchange time across the living human cortex was estimated to be about 13 ms on Connectome 2.0, faster than measured using a protocol compatible to Connectome 1.0
- Spatially varying exchange times were observed across the cortex
- A GPU-accelerated tool was developed to speed up parameter estimation by 100 times
- The narrow pulse approximation used in neurite exchange imaging does not affect estimation performance in high gradient performance MRI systems

## 1. Introduction

Understanding the dynamics of water exchange within the human brain provides a key window into deciphering pathological mechanisms and assessing tissue integrity in aging and a variety of neurological and psychiatric disorders (Bai et al., 2019, 2018). In neural tissues, water can passively traverse lipid bilayers or via aquaporins, specialized membrane protein channels that facilitate water movement between extracellular and intracellular compartments (Papadopoulos and Verkman, 2013). Disruptions in water homeostasis in conditions such as brain edema or ischemic stroke may alter membrane permeability between water compartments (King et al., 2004). Recent research has also highlighted the role of ATPase channels in actively co-transporting water for maintaining osmotic balance and supporting normal brain function (MacAulay, 2021; Springer et al., 2023b, 2023a; Williamson et al., 2023). Characterizing membrane permeability may offer new insights into physiological processes and holds promise for disease monitoring and therapeutic intervention (Badaut et al., 2014). By elucidating these mechanisms, researchers can potentially identify novel targets for intervention and refine treatment strategies aimed at preserving brain health.

Diffusion MRI (dMRI) is a versatile imaging modality for investigating brain tissue microstructure noninvasively (Basser, 1997; Le Bihan, 1995). By harnessing magnetic field gradients, dMRI encodes the displacement of water molecules through the signal attenuation incurred as water molecules diffuse randomly and cause irreversible phase dispersion (Carr and Purcell, 1954; Stejskal and Tanner, 1965). In neural tissues, water diffusion is often restricted or hindered by cell membranes and organelles (Beaulieu, 2002). Understanding how diffusion-induced signal attenuation varies as a function of diffusion-encoding gradient field strength, duration, and diffusion times with respect to tissue microscopic structure promises to yield valuable histological-level information. One of the most widely applied uses of dMRI is the study of white matter microstructure (Assaf et al., 2008, 2004; Fan et al., 2020; Fieremans et al., 2011; Huang et al., 2020; Jespersen et al., 2007; Kaden et al., 2016; Zhang et al., 2012). The prevailing biophysical model of diffusion in white matter involves a collection of fiber bundles (fascicles), each comprised of aligned anisotropic water compartments with Gaussian diffusion, characterized by distinct diffusivities and water fractions, corresponding to the water inside and outside axons (Novikov et al., 2018b, 2019). Due to the presence of myelin sheath, water exchange between the compartments within a bundle is neglected over typically used diffusion times (Nilsson et al., 2013; Veraart et al., 2020, 2019). The orientations of such multicompartmental fiber bundles within a voxel are characterized by a fiber orientation distribution function (ODF). This model, involving both the compartmental parameters of the bundle and the orientational parameters of the ODF, is commonly referred to as the Standard Model of diffusion in white matter (Novikov et al., 2018b, 2019).

Extending the Standard Model onto gray matter requires accounting for its specific microstructure properties. First, the soma compartment, corresponding to the cell bodies of neurons and glial cells, constitutes a significant portion of the gray matter volume (10-20%) (Shapson-Coe et al., 2024) and has a distinct isotropic and fully restricted contribution to the MRI signal (Palombo et al., 2020). Second, while myelin acts as a major barrier to diffusion and intercompartmental exchange in white matter, its concentration in gray matter is notably lower (Shapson-Coe et al., 2024), though both myelinated and unmyelinated axons can coexist in a voxel. The thin, branch-like cellular structures (neurites), such as unmyelinated axons, dendrites and glial cell processes, may contribute to a non-negligible water exchange between neurites and extracellular space across the neurite membrane given the relatively high surface-area-to-volume ratio and/or other water transport mechanisms, such as via aquaporins and ATPase channels (MacAulay, 2021; Papadopoulos and Verkman, 2013; Springer et al., 2023b, 2023a; Veraart et al., 2020; Williamson et al., 2023).

The presence of intercompartmental water exchange and restricted diffusion can be disentangled through multidimensional spectroscopy methods like diffusion exchange spectroscopy (DEXSY) (Callaghan and Furó, 2004). In DEXSY, water exchange between environments with different diffusivities is observed during the mixing time of a double diffusion encoding scheme, manifesting as off-diagonal elements in the two-dimensional spectrum (Callaghan and Furó, 2004). However, DEXSY demands a large amount of data to compute the multidimensional exchange spectrum, which limits its direct applicability in imaging contexts. Efforts have been made to shorten the lengthy acquisition of DEXSY (Åslund et al., 2009; Cai et al., 2018; Williamson et al., 2020, 2019), notably through filtered exchange imaging (FEXI) (Lasič et al., 2011), a variant tailored for clinical feasibility. Nonetheless, the interpretation of apparent exchange rates derived from FEXI can be confounded by water exchange between the intra- and extra-vascular environments (Bai et al., 2020) and the geometry of the involved compartments (Khateri et al., 2022; Li et al., 2023). Despite the challenges facing FEXI, previous research underscores the association between apparent exchange rates and membrane permeability (Nilsson et al., 2013; Ning et al., 2018). To shorten the lengthy multi-dimensional DEXSY-like experiments, tissue microstructure and intercompartmental water exchange can be assessed by employing a parsimonious *model* of exchange between just a few compartments. Models reduce the number of parameters making their estimation more robust and less SNR-demanding, yet they rely on explicit assumptions for the microstructure (Novikov et al., 2018a). The most widespread model of exchange in the dMRI context has been suggested by Kärger (Kärger, 1985). In its original formulation, exchange was postulated to occur between two isotropic compartments with otherwise Gaussian diffusion, at each point of the sample, governed by simple rate equations in the spirit of (Zimmerman and Brittin, 1957).

Extending the Kärger model (KM) onto structurally rich microstructure involved clarifying its assumptions. It was realized that KM applies only at sufficiently long diffusion times, after *coarse-graining* the tissue microstructure by the diffusion process, i.e., at the scales exceeding cell sizes and the correlation length *l*_*c*_of cell packing, assuming that the exchange time *t*_*ex*_ ≫ *t*_*c*_is also slow on the scale of the corresponding correlation time 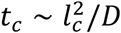 (Fieremans et al., 2010). In other words, KM is an *effective theory* valid at long times *t*, *t*_*ex*_ ≫ *t*_*c*_ and small wave vectors *q* ≪ 1/*l*_*c*_, at which point the assumptions of Gaussian diffusion and exchange occurring at every point in space can become justified, and ordinary rate equations can be used instead of solving a genuine partial differential equation for the diffusion in a complex tissue geometry.

Compartment anisotropy, essential for neuronal tissue, can be further incorporated into KM (understood as an effective theory), using the steps analogous to building the Standard Model. First, exchange within a perfectly aligned fiber bundle (fascicle) of neurites and their extra-neurite (extracellular) space is added (assuming *t*, *t*_*ex*_ ≫ *t*_*c*_), yielding the *anisotropic Kärger model* for aligned fibers (Fieremans et al., 2010). Next, as in the Standard Model, the response of such an elementary bundle is convolved with an arbitrary fiber ODF, leading to the Neurite Exchange Imaging (NEXI) (Jelescu et al., 2022) and Standard Model with Exchange (SMEX) (Olesen et al., 2022) models suggested recently for gray matter, where the parameter constraint of isotropic extra-neurite space is further employed for the estimation robustness. We can thereby refer to NEXI/SMEX as the implementations of a *locally anisotropic* (or microscopically anisotropic) Kärger model, even if the overall signal from gray matter can be fairly isotropic due to an almost isotropic ODF.

Fitting such anisotropic extensions of the Kärger model, or its cumulants such as the time-dependent kurtosis (Fieremans et al., 2010), to the diffusion signals measured at multiple diffusion times enables the quantification of exchange time scales (Jelescu et al., 2022; Lee et al., 2020; Olesen et al., 2022). A prominent indicator showing the dominance of the exchange phenomenon is the relative decrease in the direction-averaged (spherical mean) signal as diffusion time increases for the same diffusion weighting, a pattern observed in recent investigations focusing on the gray matter of the mouse (Jelescu et al., 2022; Olesen et al., 2022) and human brain (Lee et al., 2022; Uhl et al., 2024) and in stoke (Lampinen et al., 2021; Lätt et al., 2009). The main advantage of the Kärger model is its straightforward implementation using the conventional pulsed-gradient dMRI sequence, despite the demands on gradient hardware to detect fast exchange effects (Williamson et al., 2023, 2019) at short diffusion times. To facilitate efficient model fitting for exchange time evaluation, the narrow pulse approximation has been used to derive the analytical solution of the anisotropic Kärger model (Jelescu et al., 2022), though the potential impact of this approximation on exchange time measurements in clinically relevant scenarios with varying signal-to-noise ratios (SNR) remains unclear. Clarifying this aspect is crucial for the robust interpretation of exchange dynamics in clinical and preclinical settings.

Recent advances in MRI gradient systems present significant opportunities for *in vivo* tissue microstructure imaging using dMRI (Feinberg et al., 2023; Foo et al., 2020; Huang et al., 2021). The first-generation Connectome MRI scanner (Connectome 1.0; C1) was equipped with a high-performance gradient system achieving a maximum strength of 300 mT/m and a maximum slew rate of 200 T/m/s (Setsompop et al., 2013). This system enabled *in vivo* human dMRI with enhanced diffusion weighting and superior SNR compared to clinical scanners. Recent studies have demonstrated the feasibility of employing such a system to probe the *in vivo* exchange time using pulsed-gradient based NEXI (Uhl et al., 2024) or dMRI of free gradient waveform (Chakwizira et al., 2023). Recently, a second-generation Connectome MRI scanner (Connectome 2.0; C2) was developed, featuring an improved high-performance gradient system capable of generating the strongest gradients for head-only *in vivo* human imaging at 3T to date, with a maximum gradient strength of 500 mT/m and a maximum slew rate of 600 T/m/s (Huang et al., 2021). These enhancements offer the potential for dMRI measurements at even stronger diffusion weighting with reasonable SNR, facilitating the exploration of intercompartmental exchange effects across a wider range of diffusion times using the Karger models.

The increasing availability of high-performance gradient systems for *in vivo* imaging, such as the Siemens Cima.X system (maximum strength of 200 mT/m, maximum slew rate of 200 T/m/s, whole-body 3T), GE SIGNA MAGNUS system (Foo et al., 2020) with HyperG gradients (maximum strength of 300 mT/m, maximum slew rate of 750 T/m/s, head-only 3T) and high-performance gradient insert (maximum strength of 200 mT/m, maximum slew rate of 600 T/m/s) (Weiger et al., 2018), provides essential feasibility to study the intercompartmental exchange times on *in vivo* human brain using NEXI. The primary goal of this study has been to investigate intercompartmental exchange times across the living human cortex using the stronger and faster gradients of the Connectome 2.0 MRI scanner compared to those estimated with a protocol compatible with the Connectome 1.0-alike systems. Furthermore, we sought to evaluate the impact of the narrow pulse approximation on exchange time estimation. Finally, we demonstrated a novel tool leveraging GPU processing to accelerate the NEXI model fitting process, thereby enhancing the efficiency and scalability of parameter estimation for tissue microstructure imaging.

## 2. Theory

### 2.1. Anisotropic Kärger Model

We utilized the anisotropic Kärger model (Fieremans et al., 2010; Jelescu et al., 2022; Olesen et al., 2022) to model the exchange effect between neurites and extracellular water in gray matter (Figure 1a). Briefly, the anisotropic Kärger model comprises two exchanging Gaussian compartments: an anisotropic “stick”-like neurite compartment with zero diffusivity perpendicular to the long axis, and its local extracellular space. We assumed isotropic diffusion in the extracellular space to simplify the signal model and improve the robustness of estimation by reducing the number of model parameters. Considering a medium composed of neurites (n) and the extracellular water (e), its signal representation can be written as (Fieremans et al., 2010; Kärger, 1985; Ning et al., 2018):

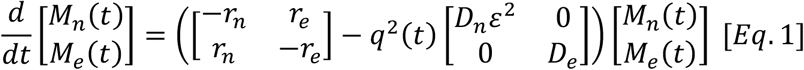

where *M*_{*n*,*e*}_(*t*) are the both T_1_-weighted and T_2_-weighted magnetizations of neurite and extracellular water, respectively, at diffusion time *t*, *r*_{*n*,*e*}_are the exchange rates from neurite to extracellular water and vice versa, such that the two compartments satisfy the detailed balance *r*_*n*_*f*_*n*_ = *r*_*e*_*f*_*e*_ with neurite volume fraction 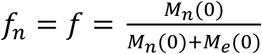 and extracellular volume fraction *f*_*e*_ = 1 − *f*, and *D*_{*n*,*e*}_ are the along-neurite and isotropic extracellular water diffusivities, respectively. Here, we assumed the longitudinal and transverse relaxation times of the neurite and extracellular water to be the same, and the echo time dependence due to transverse relaxation and the effect of coil sensitivity are cancelled out after normalized the measurements by the *b*=0 signal. The diffusion wave vector *q*(*t*) represents the diffusion gradient profile *G*(*t*) such that 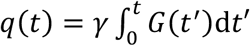 with the gyromagnetic ratio *γ* and *b*-value 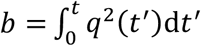 defining the diffusion weighting. The directional dependence is embodied in the dot product 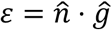 between the neurite orientation 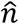 and the diffusion gradient direction 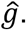 The exchange time between the neurite and extracellular water is defined as *t*_*ex*_ = 1/(*r*_*n*_ + *r*_*e*_) = (1 − *f*)/*r*_*n*_. To account for the fiber dispersion of neurites in an MRI voxel, the diffusion signal 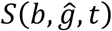 can be expressed as the convolution of the overall signal kernel *K*(*b*, *ε*, *t*) = *M*_*n*_ + *M*_*e*_ from Eq. (1) with the ODF 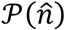 (Novikov et al., 2018b):

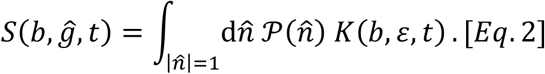

**Figure 1:**
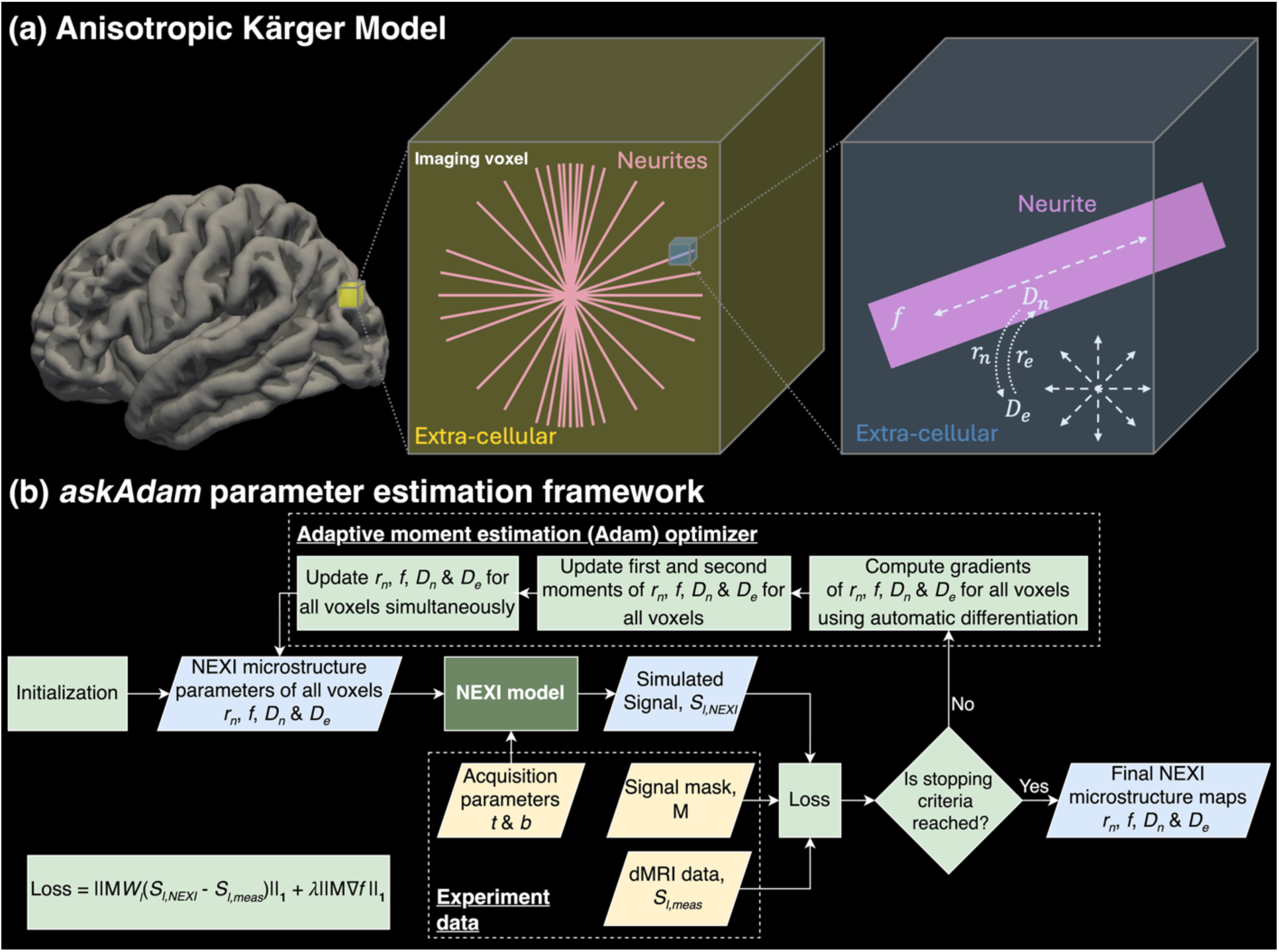
(a) Graphical illustration of the anisotropic Kärger model comprising the neurite compartment and extracellular water compartment. Each MRI voxel contains multiple neurites with various orientations. Intercompartmental water exchange takes place between neurites and extracellular water through various water movement mechanisms. (b) Illustration of the askAdam parameter estimation approach used in this work.

For any linear tensor encoding gradient waveform, e.g., pulsed-gradient sequence of finite pulse width (Stejskal and Tanner, 1965), the overall fiber response *K*(*b*, *ε*, *t*) entering Eq. (2) can be calculated by numerically solving the system of coupled differential equations Eq. (1). Here, we refer to this method as SMEX (Olesen et al., 2022). It is worth noting that while the soma compartment is not explicitly accounted for in the NEXI/SMEX model, it is considered part of the extracellular compartment by assuming the soma and extracellular water have similar diffusion properties and/or being well-mixed (Olesen et al., 2022). Further, the exchange between soma and extracellular compartments is assumed negligible due to the small surface-to-volume ratio of somas, compared with neurites. More details about the theory and assumptions can be found in (Olesen et al., 2022).

While Eq. (1) is a general representation of the fiber bundle (fascicle) response for any gradient profile, computing the dMRI signals requires solving the coupled equations in Eq. (1) using numerical solvers, which is computationally costly and results in a lengthy model fitting process. Alternatively, under the narrow pulse approximation, the solution of Eq. (1) has an analytical form (Fieremans et al., 2010; Kärger, 1985):

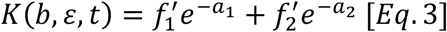

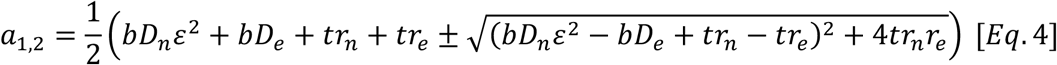

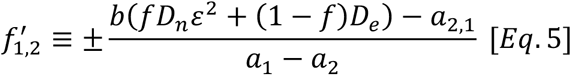

which has been referred to as NEXI (Jelescu et al., 2022).

The NEXI microstructure parameters, *f*, *D*_*n*_, *D*_*e*_ and *r*_*n*_, can be estimated using the rotationally invariant mapping framework (Novikov et al., 2018b; Reisert et al., 2017) in which both the signal model and the measured dMRI signal are projected onto the spherical harmonics space after normalizing the diffusion-weighted signal by *b*=0 signal. The *l*-th order rotationally invariant signal *S*_*l*_ can be represented as:

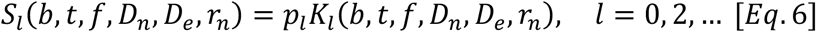

where the rotational invariant *p*_*l*_ characterizes the ODF anisotropy for each order *l* and *p*_0_ ≡ 1 (ODF normalization), and *K*_*l*_ is the projection of signal kernel *K*(*b*, *ε*, *t*) onto the Legendre polynomials. Specifically, the zeroth-order signal invariant *S*_*l*=0_ corresponds to the commonly known directionally averaged, spherical mean signal (Kaden et al., 2016). Higher-order invariants *S*_*l*=2,4,…_may also contain microstructure information when the ODF is anisotropic.

### 2.2. GPU-accelerated ‘askAdam’ framework for NEXI parameter estimation

Estimation of the NEXI microstructure parameters from Eqs. [3-6] involves optimization on a 4-dimensional parameter space (one more additional dimension for every order *l* from *l* = 2,4,6, …). Utilizing conventional nonlinear least-square approaches based on CPU computation for NEXI parameter estimation can be computationally lengthy for whole-brain mapping: the fitting process is repeated hundreds of thousands of times sequentially over all voxels in an imaging volume. Recent research has demonstrated the efficiency of accelerating the parameter estimation process based on GPU computation, both on minimization based (Harms et al., 2017; Hernandez-Fernandez et al., 2019) and supervised learning-based approaches (Jung et al., 2021; J. Lee et al., 2019; Martins et al., 2021; Nedjati-Gilani et al., 2017). Here, we incorporated NEXI parameter estimation into the ‘askAdam’ framework (Chan et al., 2022; Marques et al., 2023), one of the minimization-based GPU-accelerated methods, aiming to provide an additional resource to shorten the estimation processing time while maintaining similar estimation performance to the conventional CPU-based non-linear least-square (NLLS) approach.

‘AskAdam’ is a parameter estimation tool (Chan et al., 2022; Marques et al., 2023), leveraging the efficiency and suitability of the stochastic gradient-descent-based optimizer, such as Adam, to handle large non-convex problems. The Adam optimizer is commonly used in deep learning to perform learnable network parameter updates (Kingma and Ba, 2014) (Figure 1b). To enable askAdam to process multiple voxels at the same time in NEXI parameter mapping, the optimization problem can be formulated as:

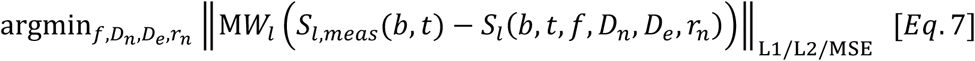

where M is the (brain) signal mask, 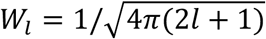 is the weight associated with the *l*-th order rotationally invariant signal (Novikov et al., 2018b) and *S*_*l*,*meas*_ is the *l*-th order rotationally invariant signal from the actual measurements. The optimization loss shown in Eq. (7) is nearly identical to that used for an NLLS solver in conventional voxelwise fitting. The main difference is that all input voxels inside the signal mask (which can be a full set or subset of the whole-brain data, depending on the memory allowance of the GPU) contribute to one loss in a single optimization problem in the askAdam framework. Here, the initialization of the NEXI microstructure parameters can be done by using random values or prior estimates as in NLLS. During each optimization cycle, the NEXI microstructure parameters of all voxels are incorporated with the sequence settings (e.g., diffusion times and *b*-values) to forward simulate the NEXI signals across the entire volume using elementwise arithmetic operations to derive the loss in Eq. (7). The loss gradient with respect to the model parameters on each voxel is then computed using the automatic differentiation function from Matlab, so that the model parameter across the entire dataset can be updated simultaneously using the Adam optimizer (Figure 1b). All computations (i.e., forward signal generation, gradient computation and parameter update) can take place on a GPU to accelerate the optimization process. This process is repeated until the stopping criteria are fulfilled (either the maximum number of optimization iterations is reached or the change of loss across iterations is less than a certain tolerance value; a maximum of 4000 iterations and a tolerance level of 1×10^-8^ were chosen here). We choose the L1-norm as the loss function in Eq. (7) as it outperforms the L2-norm at moderate SNR in our preliminary analysis (see Figure S1).

Another advantage of askAdam is that the loss function can be further customized to include spatial regularization as part of the optimization problem, which has been used in image reconstruction (Knoll et al., 2011), quantitative susceptibility mapping (QSM Consensus Organization Committee et al., 2024) and multi-compartment parameter fitting (Orton et al., 2014; Pasternak et al., 2008) to improve the estimation results. To do so, the loss function of Eq. (7) can be generalized into:

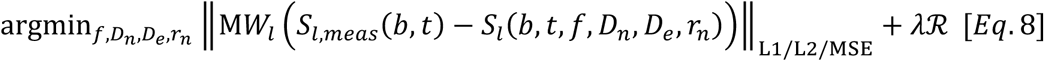

where *λ* is a regularization parameter and ℛ can be any regularizer that promotes certain features (for example, sparsity or smoothness) in the estimation maps. This feature is feasible for askAdam because the microstructure parameters of all voxels update simultaneously during the model fitting process, in turn, allowing the regularizer to stay up to date during the optimization cycle.

## 3. Methods

### 3.1. Numerical simulations

#### 3.1.1 Noise propagation with forward SMEX and NEXI signal generation

To address the research question of whether the narrow pulse solution employed in NEXI is sufficient to represent the dMRI signal in an actual experiment for high-gradient performance MRI systems, we conducted a noise propagation analysis to evaluate the estimation performance of the NEXI model fitting using SMEX and NEXI for forward signal generation with realistic dMRI acquisition protocols designed for either the Connectome 1.0 or Connectome 2.0 scanner as shown in Table 1. We randomly chose 10,000 different combinations of the NEXI microstructure parameters within the following range: *t*_*ex*_=[1, 50] ms, *f*=[0.01, 0.99], *D*_*n*_=[0.1, 3] μm^2^/ms, and *D*_*e*_=[0.1, 3] μm^2^/ms with *D*_*n*_≥ *D*_*e*_. Complex-valued Gaussian noise was applied to the simulated signals to simulate a clinically relevant SNR of 50 in the non-diffusion-weighted *b*=0 signal. Realistic Rician noise conditions were simulated by extracting the magnitude of the signal with complex-valued Gaussian noise. NEXI model fitting was subsequently performed on the spherical mean (zeroth order rotational invariant) after normalized to *b*=0 images. The starting point of the estimation for each voxel was set up using a dictionary-matching approach (Ma et al., 2013), achieved by searching the maximum likelihood of the inner product of the *in vivo* measured signal to a dictionary of 10,000 randomly generated simulated signal across a predefined range of *r*_*n*_=[0.001, 0.99] s^-1^, *f*=[0.01, 0.99], *D*_*n*_=[1.5, 3] μm^2^/ms and *D*_*e*_=[0.5, 1.5] μm^2^/ms. The fitting boundary of each NEXI parameter was set to: *r*_*n*_=[0, 1] s^-1^, *f*=[0, 1], *D*_*n*_=[0.1, 3] μm^2^/ms, *D*_*e*_=[0.1, 3] μm^2^/ms, and we did not impose the same constrain of *D*_*n*_≥ *D*_*e*_as in the ground truth used in forward signal simulation. The forward SMEX and NEXI signals were calculated for 64 diffusion gradient directions per shell. Additionally, we investigated a noise-floor-correction method for the signal with Rician noise by integrating the Rician noise floor within the signal model by using the Rician mean model (Jelescu et al., 2022). We repeated this process based on the Connectome 2.0 protocol with varying diffusion gradient pulse duration *δ* = 8 ms and 10 ms, respectively, to consider its impact on Connectome 1.0-alike systems with a relatively lower maximum gradient amplitude and slower gradient slew rate.

**Table 1:** Data acquisition protocols for dMRI in this study. The same protocol parameters are used in the noise propagation analysis and *in vivo* imaging. Diffusion gradient duration is defined as the full width at half maximum of the gradient pulse. The maximum ramp time of the gradient pulse was (500/600=)0.83 ms for the Connectome 2.0 protocol and (300/200=)1.5 ms for the Connectome 1.0 protocol. For each diffusion time, the gradient strength was the only control variable to achieve various b-values, and the highest b-value utilized the maximum gradient strength of 300 mT/m on the Connectome 1.0 protocol and 500 mT/m on the Connectome 2.0 protocol. \**b*=1 ms/μm^2^ data was not used in data fitting.

| Protocol | Connectome 2.0 |  |  | Connectome 1.0 |  |  |
| --- | --- | --- | --- | --- | --- | --- |
| Diffusion time $t$ (ms) | 13 | 21 | 30 | 21 | 30 | 40 |
| Diffusion gradient duration $\delta$ (ms) | 6 | | | 10 | | |
| # gradient directions per shell | 32 (for C1 vs C2 comparison)/64 (for high SNR cortical investigation) |  |  | 32 |  |  |
| # shells | 5 | 6 | 7 | 5 | 6 | 7 |
| b-values (ms/$\mu\text{m}^2$) | [1*,2.3,3.5,4.8,6.5] | [1*,2.3,3.5,4.8,6.5,11.5] | [1*,2.3,3.5,4.8,6.5,11.5,17.5] | [1*,2.3,3.5,4.8,6.5] | [1*,2.3,3.5,4.8,6.5,11.5] | [1*,2.3,3.5,4.8,6.5,11.5,17.5] |
| T<sub>acq</sub> (min) | 21 | 25 | 29.5 | 24 | 29 | 34 |

#### 3.1.2. Noise propagation with *l*_*max*_ = 0 and *l*_*max*_ = 2 NEXI models

As introduced in the Theory section, NEXI model fitting is not limited to the spherical mean signal (zeroth order rotational invariant), but can involve any spherical harmonics order up to *l*_*max*_, depending on the total number of unique gradient directions acquired for each *b*-value. In the second noise propagation analysis, we investigated if the estimation performance could be improved when the second-order rotational invariants of the dMRI signals were added to the model fitting. From now on, we refer to fitting the zeroth order rotational invariant *S*_*l*=0_ of NEXI as *l*_*max*_ = 0, and to fitting both invariants ( *S*_*l*=0_ + *S*_*l*=2_) of NEXI as *l*_*max*_ = 2. Noise propagation was conducted under the same conditions as in Section 3.1.1, and the range of the non-linear neurite dispersion index *p*_2_ was within [0, 0.3] for the forward simulations. The *p*_2_ value increases with the tissue anisotropy (Novikov et al., 2018b). The low *p*_2_ value in simulations was consistent with the low anisotropy observed in gray matter.

#### 3.1.3. Comparison between NLLS and askAdam on NEXI parameter mapping

The askAdam framework allows data fitting to be performed simultaneously in multiple spatial dimensions with the option to incorporate spatial regularization, which is distinct from voxel-wise NLLS fitting. Therefore, we wanted to understand how this spatial regularization would affect the estimation performance between the two solvers via an *in silico* head phantom experiment. The phantom was created using the *in vivo* NEXI fitting results of one participant scanned with the Connectome 2.0 scanner (see details in Section 3.2.4). The median value across each brain parcellation label (excluded ventricles and cerebrospinal fluid) was computed for all NEXI microstructure parameters and used as the ground truth for the corresponding region of interest (ROI) segmented using FreeSurfer (Section 3.2.2). We then forward simulated the NEXI signals using the Connectome 2.0 dMRI acquisition protocol in Table 1. Gaussian noise was added to the simulated signals so that we could investigate the estimation performance at a moderate SNR of 50 and a low SNR of 25. Tissue microstructure parametric maps were derived using the NEXI *l*_*max*_ = 2 model implementation via either NLLS or askAdam solvers. Additionally, we incorporated anisotropic, two-dimensional spatial total variation (TV) regularization on the neurite fraction map for the NEXI model fitting with askAdam (askAdam_TV_). This was achieved by applying a 2D gradient operator (∇= [∇_x_; ∇_y_]) on the *f* map in the in-plane direction, i.e., ℛ = ‖M∇*f*‖_1_ in Eq. (8) (Figure 1b). The regularization parameter *λ* is empirically set to 0.002 mm (see Supplementary Figure S2 for details). We evaluated the accuracy of estimation via the median differences (bias) between the estimated values and ground truth, and the precision via the interquartile range (IQR) of the fitted values within each ROI. For all data fitting, the NLLS approach was performed on an EPYC 7313 16-Core Processor (AMD, Santa Clara, US), whereas the askAdam approach was performed on an A40 GPU (NVIDIA, Santa Clara, US).

### 3.2. *In vivo* MRI

#### 3.2.1 Data Acquisition

##### Comparison between Connectome 1.0 and Connectome 2.0 protocols

Data acquisition was performed on the 3T Connectome 2.0 scanner (MAGNETOM Connectom.X, Siemens Healthineers, Erlangen, Germany) equipped with a maximum gradient strength of 500 mT/m and slew rate up to 600 T/m/s. Imaging experiments were performed on 5 healthy volunteers (mean±SD age = 26±6 years; 3 females, 2 males) using a custom-built 72-channel head coil for signal reception (Mahmutovic et al., 2024). The study was approved by the local ethics committee, and written informed consent was obtained from all participants for being included in the study. The imaging protocol comprised:

1. Whole-brain T_1_-weighted scan using MPRAGE with 0.9 mm isotropic resolution, acquisition time = 5.5 min;
2. 2D spin-echo EPI-dMRI, multi-band factor of 2, GRAPPA factor of 2, partial Fourier of 6/8, resolution = 2 mm isotropic, TR/TE_C1_/TE_C2_ = 3600/68/54 ms using the diffusion scheme laid out in Table 1. Non-diffusion-weighted images (*b*=0) were acquired interspersed for every 16 diffusion-weighted images (DWIs). Total acquisition time = 40 min per protocol.

##### NEXI measurements with higher SNR data

Additionally, 10 healthy volunteers (mean±SD age = 31±15 years; 6 females, 4 males) were scanned using the same hardware and imaging sequences to measure the NEXI’s exchange time with higher SNR data. The same C2 protocol as described in Table 1 was used for this dataset, with the only modification being that 64 diffusion encoding directions were acquired for each b-value instead of 32. The total acquisition time is 80 min.

#### 3.2.2. Data processing

The MPRAGE images were processed using FreeSurfer’s *‘recon-all’* function for cortical surface reconstruction (Fischl, 2012) and SynthSeg (Billot et al., 2023) for gray matter parcellation. Diffusion MRI data were processed based on a modified DESIGNER pipeline (Ades-Aron et al., 2018), including Rician-noise-floor-corrected Marchenko-Pastur Principal Component Analysis (MP-PCA) image denoising with 3 iterations (Tournier et al., 2023; Veraart et al., 2016), Gibbs ringing artifact removal on non-diffusion-weighted images based on local sub-voxel-shifts in MRtrix (Kellner et al., 2016; Tournier et al., 2019), susceptibility-induced image distortion using FSL’s *topup* (Andersson et al., 2003), eddy current-induced distortion using FSL’s *eddy* (Andersson and Sotiropoulos, 2016), and gradient non-linearity distortion correction with in-house code (Fan et al., 2016; Jovicich et al., 2006). Image registration between MPRAGE and dMRI data was performed based on non-linear registration using ANTs (Avants et al., 2011). The transformation matrices and warp fields were subsequently used to transform MPRAGE-derived gray matter parcellation labels to dMRI space for statistical analysis and dMRI-derived parameter maps to MPRAGE space for visualization.

#### 3.2.3. Investigating time dependence via ROI fitting

We evaluated the time dependence of the *in vivo* dMRI signals in various areas of the gray matter on ROI-averaged diffusion signals. Six ROIs, including the amygdala, hippocampus, frontal, parietal, temporal and occipital gray matter, were created by combining the parcellation labels from SynthSeg according to (Klein and Tourville, 2012). Zeroth and second-order rotationally invariant dMRI signals (*S*_*l*=0_ and *S*_*l*=2_) on each voxel were derived by projecting the directionally dependent dMRI signal up to the fourth-order spherical harmonic after normalizing the DWIs by the mean *b*=0 images (*S*_*l*=4_ was not used). The rotationally invariant signals were then averaged over each ROI and across all subjects. The averaged signals were fitted to both zeroth and second order ( *l*_*max*_ = 2) spherical harmonic of the NEXI model using the NLLS implementation. All diffusion times and *b*-values were used in the model fitting except data acquired at *b*=1 ms/μm^2^ since the restricted diffusion inside the soma compartment could still have non-negligible contributions to the dMRI signal at low *b*-values (see Supplementary Figure S6).

#### 3.2.4. NEXI microstructure parameter mapping

Tissue microstructure parametric maps were derived using the NEXI *l*_*max*_ = 2 model implementation on the whole-brain data for individual subjects via either the NLLS or askAdam solvers similar to the *in silico* head phantom experiment in Section 3.1.3. Additionally, we incorporated 2D-TV regularization for the NEXI model fitting with askAdam to validate the *in silico* experiment findings with the *in vivo* data. Subsequently, the exchange time maps were transformed into FreeSurfer’s fsaverage subject space and projected onto the cortical surface for visualization.

#### 3.2.5. Statistical analysis

In subsequent statistical analyses on the whole-brain parameter maps, the median value within each cortical ROI was computed to minimize outliers due to potential partial volume effects and imperfect cortical surface registration between the MPRAGE and dMRI images for all the NEXI microstructure parameters on each subject. Group-level statistics were derived by computing the mean exchange time and its standard deviation (SD) across subjects for each ROI. With the 2 mm isotropic resolution of the *in vivo* data, the partial volume effect originating from the cerebrospinal fluid (CSF) and superficial white matter may affect the accuracy of the estimated gray matter exchange time. Therefore, we performed a Pearson’s correlation analysis to investigate whether the exchange time derived from NEXI was associated with the cortical thickness estimated from the SynthSeg parcellation result on all ROIs. Additionally, we computed the Pearson’s correlation coefficients between the NEXI microstructure parameters to understand how they interacted with one another.

## 4. Results

### 4.1. Noise propagation analysis

Noise propagation results from fitting the zeroth order rotational invariant of NEXI to either SMEX or NEXI for forward signal generation showed similar estimation performance on all parameters at an SNR of 50 based on Connectome 2.0 dMRI acquisition protocol (Figure 2a). Similar observations can be made when the gradient pulse duration was set to 8 ms and 10 ms (Figure 2b). We compared the three noise correction scenarios: Rician uncorrected noise (RUC), Rician mean model (RM), and Gaussian noise. Among the three cases, the uncorrected Rician noise signal had the worst performance with greater biases and wider IQRs for all parameters (Figure 2c). Incorporating the Rician mean into the NEXI signal model (RM) or with the Gaussian noise greatly reduces the estimation biases and IQRs for all fitted parameters. Notably, the exchange time estimation from the Connectome 1.0 protocol is less accurate and less precise than those derived from Connectome 2.0 at faster exchange times (≤20 ms) in the presence of the Rician noise floor. In the relatively slower exchange time regime (>20 ms), the estimation performance between the C1 and C2 is comparable in all cases (left panel, Figure 2c). Comparable noise performance was observed when using only the zeroth order rotational invariant of the signal and incorporating the second order rotational invariant into the NEXI model fitting, for all estimated parameters (Figure 2d). The noise propagation analysis result shows that the additional fitting parameter *p*_2_ (corresponding to the nonlinear fiber dispersion index) can be robustly measured without degrading the estimation performance of the other parameters (Figure 2d).

**Figure 2:**
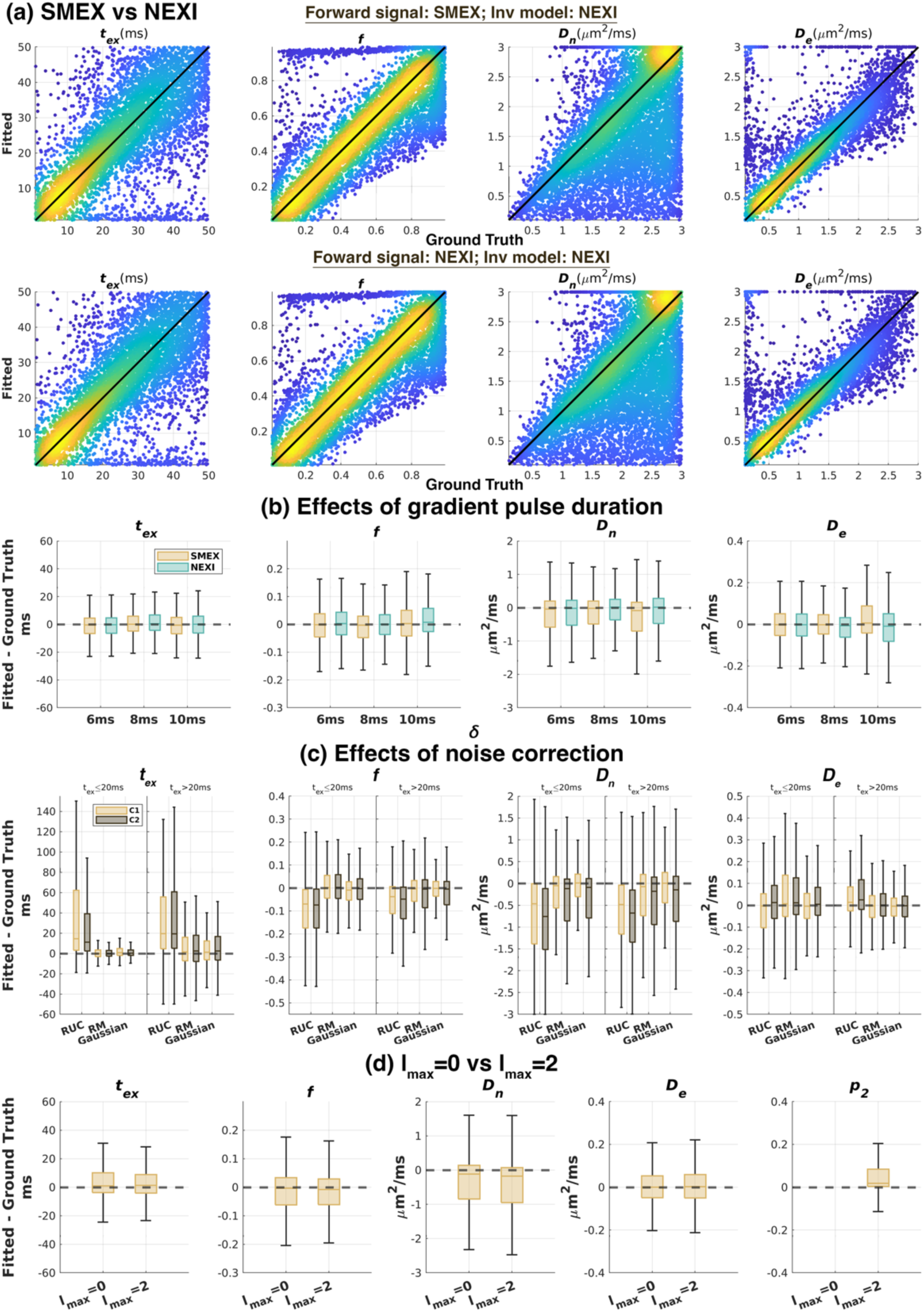
Noise propagation analysis evaluating the effect of the narrow pulse approximation for NEXI at SNR of 50. (a) Scatter plots of the estimated NEXI microstructure parameters versus ground truth values on data with Gaussian noise using Connectome 2.0 protocol. Color scale corresponds to the occurrence on the spatial grid (Yellow: high occurrence; Blue: low occurrence). (b) Box plots of the NEXI parameters estimated with various gradient pulse durations. (c) Box plots of the NEXI parameters estimated with various Rician noise floor correction methods. (d) Box plots of the NEXI parameters estimated with l_max_=0 and l_max_=2.

### 4.2. NLLS vs askAdam

To determine if askAdam can be an effective optimization tool for NEXI model fitting, we compared its estimation performance to the standard NLLS approach based on the summary statistics across ROIs using image-based numerical simulations. The processing time on the simulated whole-brain dataset phantom for the NLLS approach was about 13 hours on 1 CPU, whereas the askAdam-based approaches took about 6 min, achieving over 100-fold acceleration on processing time by GPU. In terms of accuracy and precision, we observed that askAdam was slightly less accurate than NLLS for the exchange time estimation (median bias of NLLS=0.26 ms vs askAdam=0.67 ms at SNR=50; Figure 3b-c), while the precision is comparable (median IQR of NLLS=3.01 ms vs askAdam=3.24 ms, Figure 3d). Interestingly, the measurement accuracy remains comparable with precision being improved when we introduced a weak spatial regularization with askAdam in NEXI model fitting (median bias of askAdam=0.67 ms vs askAdam_TV_=0.56 ms; median IQR of askAdam=3.24 ms vs askAdam_TV_=2.64 ms at SNR=50). As SNR decreases, both the accuracy and precision of the estimation are reduced for all solvers (Figure 3c-d).

**Figure 3:**
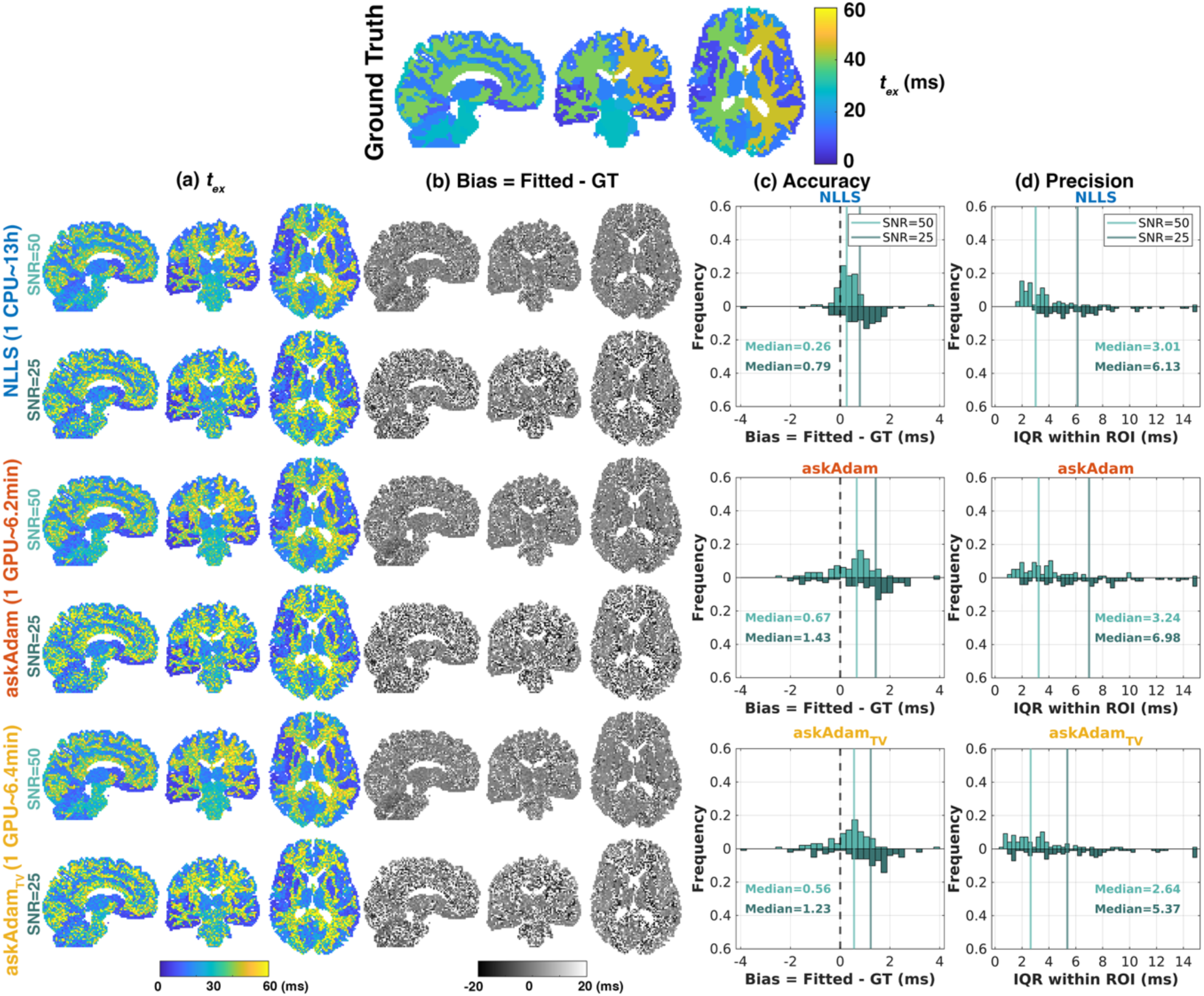
*In silico* head phantom comparison between NLLS and askAdam for NEXI parameter estimation performance. (a) The intercompartmental exchange time maps and (b) the corresponding estimation bias derived at SNR of 50 and SNR of 25 for the three solers: voxel-wise NLLS, askAdam without spatial regularization and askAdam with spatial regularization. (c) Estimation accuracy and (d) precision across all ROIs.

### 4.3. *In vivo* results

We evaluated the goodness of fit between the NEXI model fitting and the *in vivo* data using the ROI-averaged rotationally invariant signals. Time-dependent dMRI signals were observed for all gray matter ROIs (circles in Figure 4) on both Connectome 2.0 (Figure 4a) and Connectome 1.0 (Figure 4b) protocol data, where both the zeroth and second-order rotational invariants *S*_*l*=0_ and *S*_*l*=2_ decreased with diffusion time at each *b*-value. In terms of fit quality, the discrepancies between the fitted signal curves and the actual data were more noticeable on the second-order rotational invariants *S*_*l*=2_, with the amygdala showing the poorest fit in both cases, though the root mean square values of the fitting residual are comparable between *S*_*l*=0_ and *S*_*l*=2_, ranging from 2.26×10^-3^ to 3.12×10^-3^ for *S*_*l*=0_ and from 0.25×10^-3^ to 1.41×10^-3^ for *S*_*l*=2_ across ROIs. In this analysis, the estimated *t*_*ex*_ ranged from 14.93 ms (frontal) to 33.51 ms (occipital) on the Connectome 2.0 data (Table 2), whereas the estimated *t*_*ex*_ from Connectome 1.0 was substantially longer when compared to Connectome 2.0, though similar regional trends could be observed (faster exchange in the frontal, parietal and temporal gray matter).

**Figure 4:**
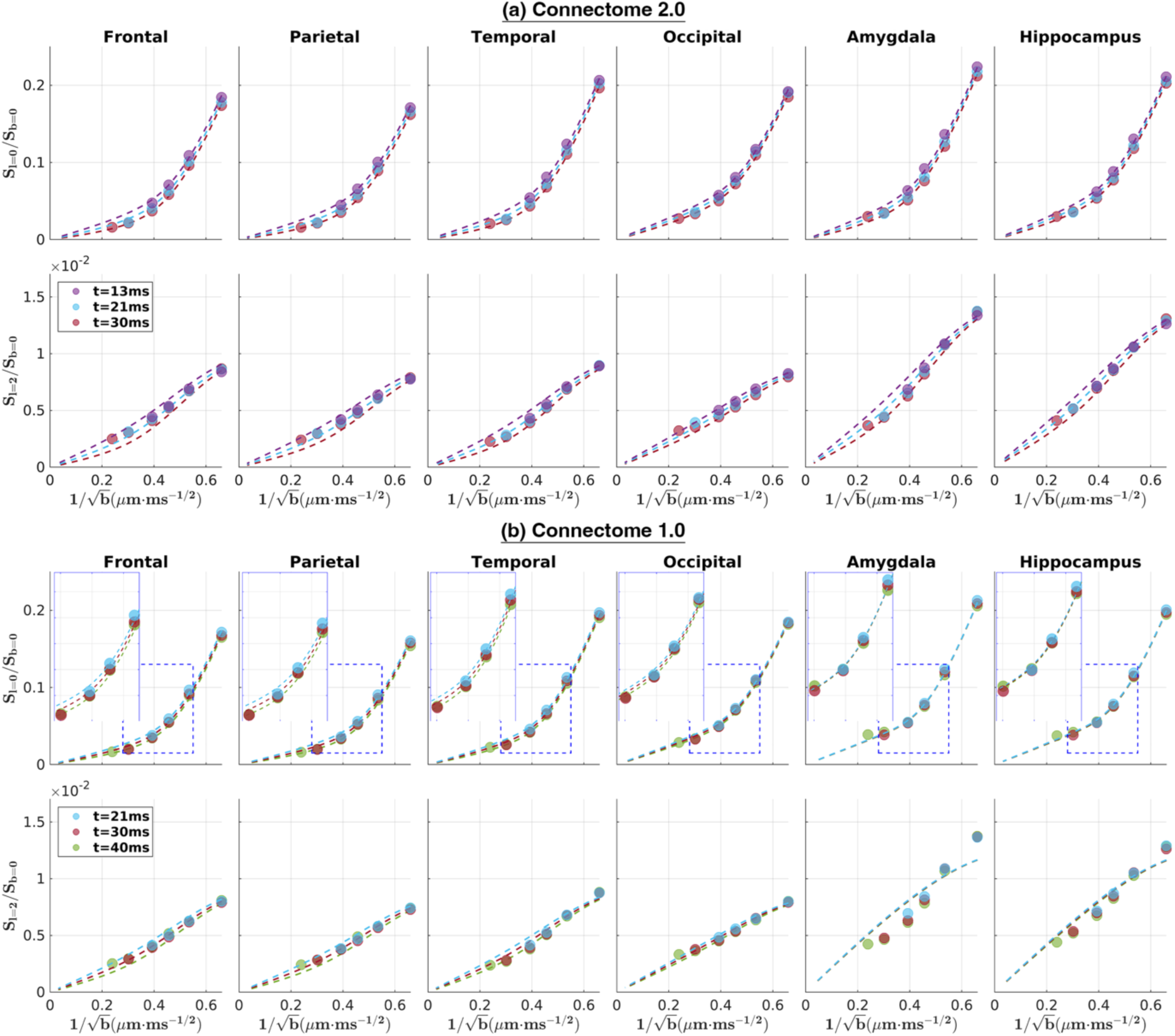
Results of ordinary NEXI model fitting on ROI-averaged *in vivo* dMRI signals. The NEXI model of *l*_*max*_ = 2 was fitted to data using the (a) Connectome 2.0, and (b) Connectome 1.0 protocol data. Circles represent the measured data and dashed lines represent the NEXI model fitting curves derived from the estimated tissue parameters.

**Table 2:** Ordinary NEXI microstructure parameters of gray matter ROI derived from ROI-averaged *in vivo* dMRI signals. C2: Connectome 2.0 protocol; C1: Connectome 1.0 protocol.

| | $t_{ex}$ (ms) | | $f$ | | $D_n$ ($\mu\text{m}^2/\text{ms}$) | | $D_e$ ($\mu\text{m}^2/\text{ms}$) | | $p_2$ | |
| --- | --- | --- | --- | --- | --- | --- | --- | --- | --- | --- |
|  | C2 | C1 | C2 | C1 | C2 | C1 | C2 | C1 | C2 | C1 |
| Frontal | 14.93 | 30.22 | 0.35 | 0.27 | 3.00 | 3.00 | 0.89 | 0.90 | 0.21 | 0.24 |
| Parietal | 17.84 | 32.15 | 0.32 | 0.27 | 3.00 | 3.00 | 0.94 | 0.95 | 0.21 | 0.23 |
| Temporal | 17.06 | 39.39 | 0.35 | 0.28 | 3.00 | 3.00 | 0.81 | 0.81 | 0.20 | 0.23 |
| Occipital | 33.51 | 77.64 | 0.34 | 0.30 | 3.00 | 3.00 | 0.89 | 0.88 | 0.19 | 0.21 |
| Amygdala | 22.96 | 605.48 | 0.37 | 0.27 | 3.00 | 3.00 | 0.77 | 0.79 | 0.28 | 0.33 |
| Hippocampus | 29.36 | 323.18 | 0.36 | 0.28 | 3.00 | 3.00 | 0.82 | 0.83 | 0.28 | 0.32 |

The *in vivo* whole-brain mapping estimation differences between the Connectome 2.0 and Connectome 1.0 protocols show similar patterns as fitting the ROI-averaged signal. The estimated *t*_*ex*_with the Connectome 1.0 protocol was about 2.67 times longer than those with the Connectome 2.0 protocol when fitting the ordinary NEXI model to the denoise dMRI data (Figure 5). Interestingly, the bias between the two protocols was reduced to a factor of 1.54 using the NEXI Rician mean model with no denoising applied to the dMRI data. For the Connectome 1.0 protocol, the *t*_*ex*_estimation of using NEXI Rician mean model was substantially shorter than those of using the ordinary NEXI model, while the *t*_*ex*_ estimations of the Connectome 2.0 protocol are comparable between the two approaches.

**Figure 5:**
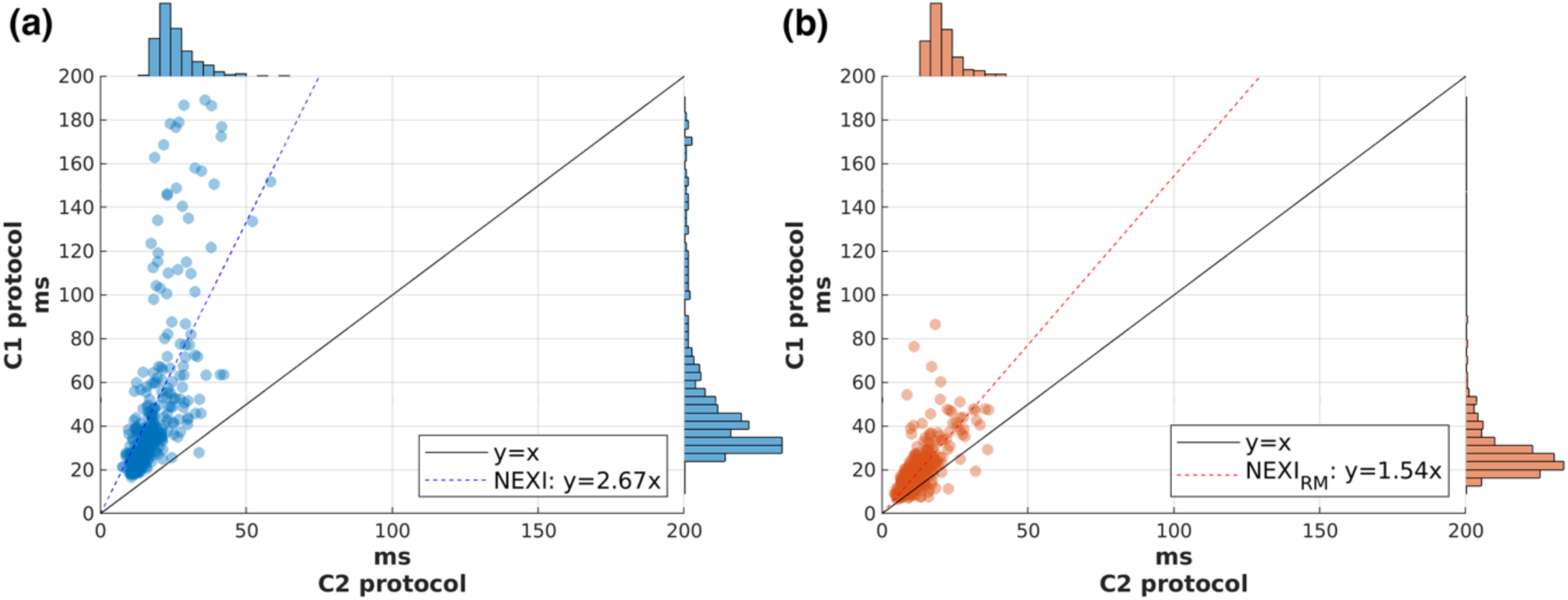
Scatterplots of the estimated exchange time using C1 and C2 protocols with (a) the ordinary NEXI model with denoise dMRI data and (b) the NEXI with Rician mean model with dMRI data that did not incorporate denoising in the preprocessing. Each data point corresponds to the median value of an ROI.

Images of both the zeroth order and second order rotational invariants of the dMRI signal demonstrated that the Connectome 2.0 scanner produced very high-quality data even at high *b*-values (Figure 6a). The mean SNR across all subjects in the cortex was 55 at *b*=0 without data denoising. The SNR of the spherical mean signal (spherical mean signal divided by the noise estimation from *b*=0 images) at *t*=30 ms for each *b*-values from 2.3 ms/µm^2^ to 17.5 ms/µm^2^ were 9.6, 5.4, 3.4, 2.4, 1.7 and 1.5, respectively. These levels can be interpreted as the ratio of spherical mean signal to the noise floor for each b-shell. On top of that, the noise fluctuation in spherical mean signal was even smaller by a factor of 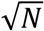 with *N* = 64 gradient directions per b-shell. The *in vivo* NEXI parameter maps on a single subject are shown in Figure 6b. Among all NEXI parameters, the intra-neurite diffusivity *D*_*n*_was the noisiest measurement. The estimated values were very close to the upper bound of the parameter range (3 μm^2^/ms), especially in white matter, whereas the estimations of neurite volume fraction *f* and ODF *p*_2_were the most robust. All three optimization solvers (NLLS, askAdam and askAdam_TV_) produced similar contrast on all parameters except *D*_*n*_, where the values from askAdam were lower than those from NLLS and close to 2.5 μm^2^/ms.

**Figure 6:**
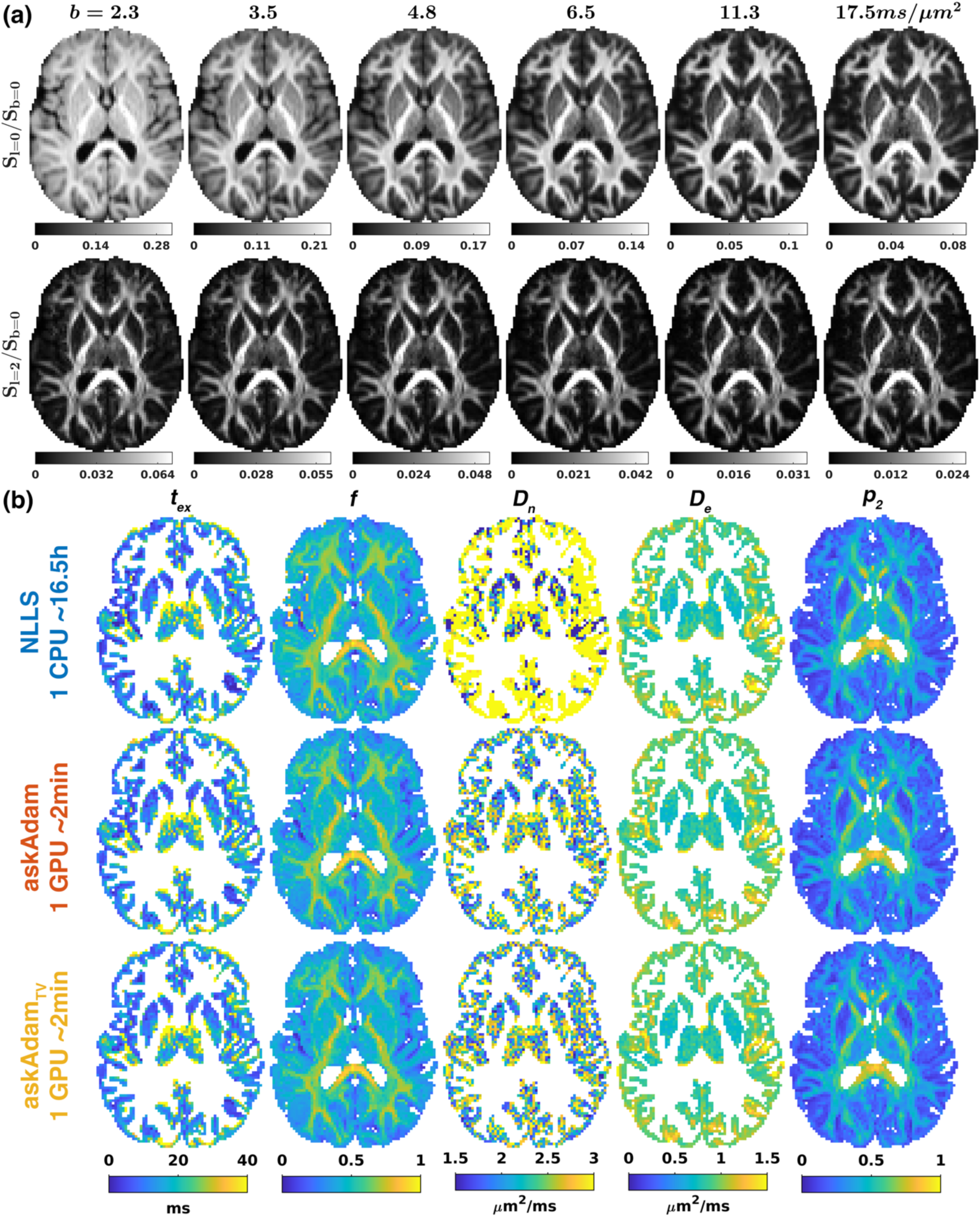
(a) Examples of the zeroth and second order rotationally invariant dMRI signals at *t*=30 ms and *b*-values=2.3-17.5 ms/µm^2^ on one subject from the high SNR Connectome 2.0 cohort. Zoom-in images can be found in Supplementary Figure S8. (b) NEXI microstructure parameters on the same subject derived from different solvers. CSF (f and p_2_) and white matter (t_ex_, D_n_ and D_e_) were masked out to provide a better illustration of the estimations on gray matter.

The exchange time maps projected onto the cortical surface of two subjects (Subject #1: 23-year-old female; Subject #2: 22-year-old male) are shown in Figure 7. In both cases, the *t*_*ex*_maps derived from NLLS were apparently noisier than those from askAdam, and the results were comparable between with and without spatial regularization for askAdam. In these two examples, both subjects showed similar spatial contrasts between the pre-/post-central gyri, visual cortex, and cingulate gyrus to their surrounding tissue.

**Figure 7:**
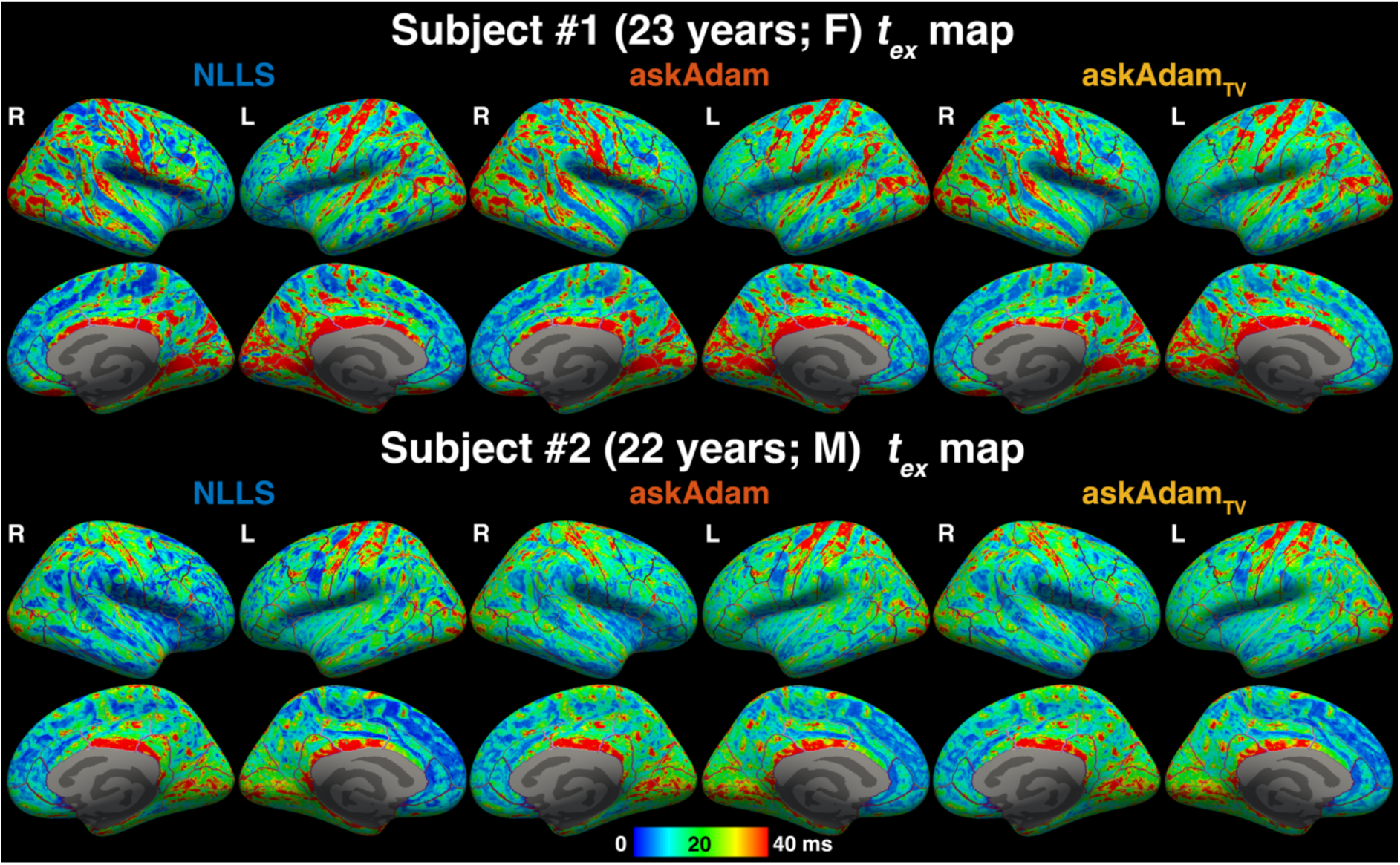
Examples of exchange time *t*_*ex*_ maps projected at middle cortical depth, halfway between the pial surface and gray-white matter interface in two subjects (Subject #1: 23 years; female; Subject #2: 22 years; male). The outline on the cortical surface indicates the boundaries of the SynthSeg parcellation labels.

In the group-level analysis, the median±IQR exchange times estimated using NLLS, askAdam, and askAdam_TV_ were 13.1±8 ms, 13.3±7 ms and 13.2±6.9 ms across all cortical ROIs for all subjects on the high SNR Connectome 2.0 data (Figure 8a). Like the spatial patterns observed at the individual level, the frontal pole, pre-/post-central gyri, visual cortex and posterior cingulate cortex had relatively longer exchange times than other ROIs on Connectome 2.0 (Figure 8b). A spatial gradient of longer exchange times was found moving from the anterior to the posterior cingulate cortex (Figure 8b-c). The mean exchange times and standard deviations across subjects were similar for all three solvers, except for the fontal pole, entorhinal, temporal pole, and parahippocampal gyrus, where the NLLS solver estimated generally longer exchange times and showed larger standard deviations (Figure 8c).

**Figure 8:**
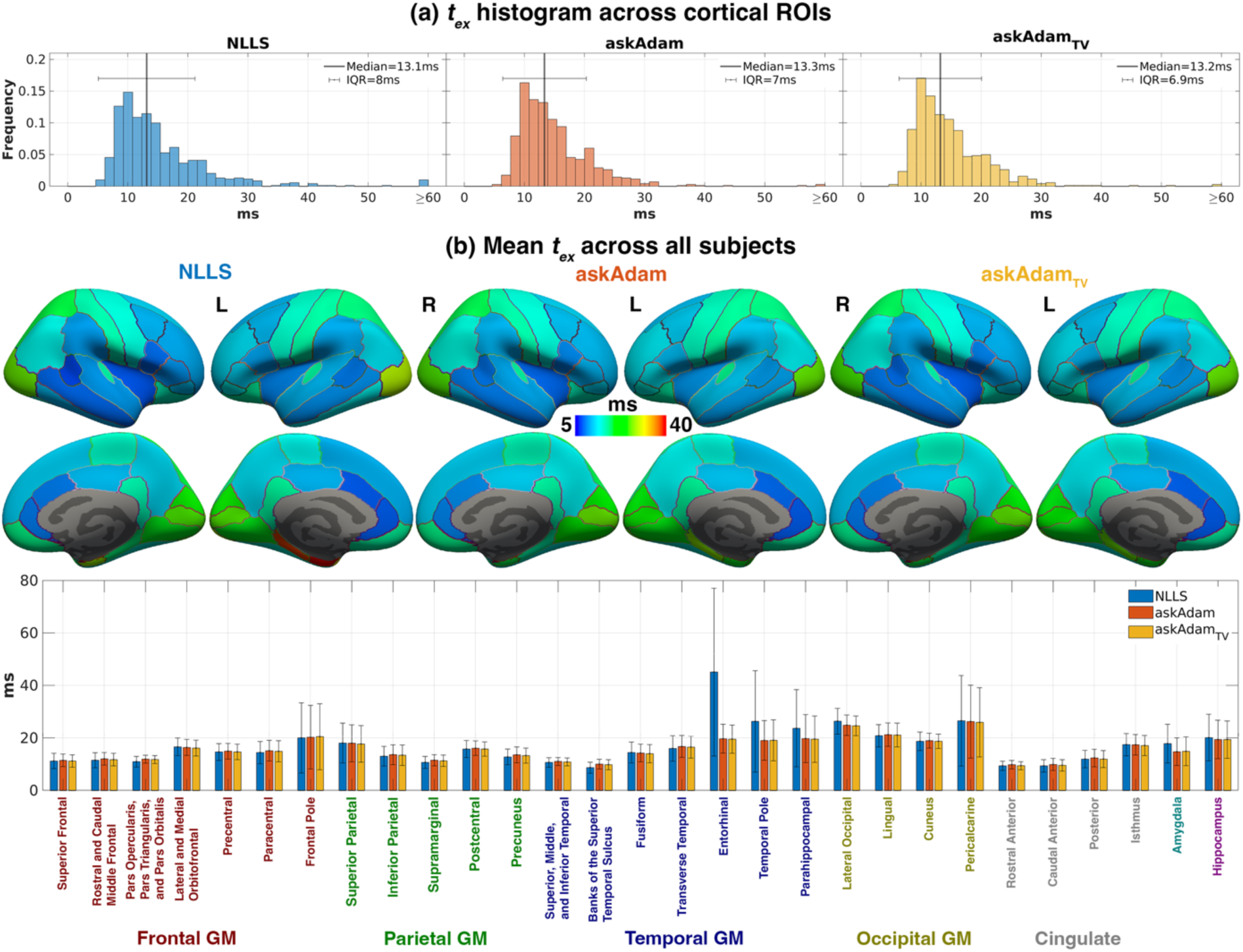
(a) Exchange time *t*_*ex*_across all cortical ROIs and all subjects using the high SNR Connectome 2.0 dataset. Solid line indicates the median exchange time of the corresponding histogram. (b) Mean and standard deviation of the exchange time across all subjects for each cortical ROIs and their corresponding projection on the cortical surface.

To investigate potential partial volume effects from CSF and superficial white matter on NEXI model fitting, we performed a Pearson’s correlation analysis between the estimated exchange time and cortical thickness for all cortical ROIs and all subjects. No significant correlations were found between these two parameters on all solvers (R=-0.01; R=-0.02 and R=-0.02 for NLLS, askAdam, and askAdam_TV_, Figure 9a). Among the NEXI tissue parameters, moderate correlations were found between *t*_*ex*_and *f* (-0.48, -0.50, -0.50), between *t*_*ex*_ and *D*_*n*_ (-0.45, -0.43) for the askAdam solvers, and between *p*_2_ and *f* (-0.53, -0.34, -0.35) for NLLS, askAdam and askAdam_TV_. Additionally, moderate correlations were observed between *p*_2_ and *D*_*n*_ (-0.37) for NLLS (Figure 9b), whereas weaker correlations between these two parameters (-0.28,

**Figure 9:**
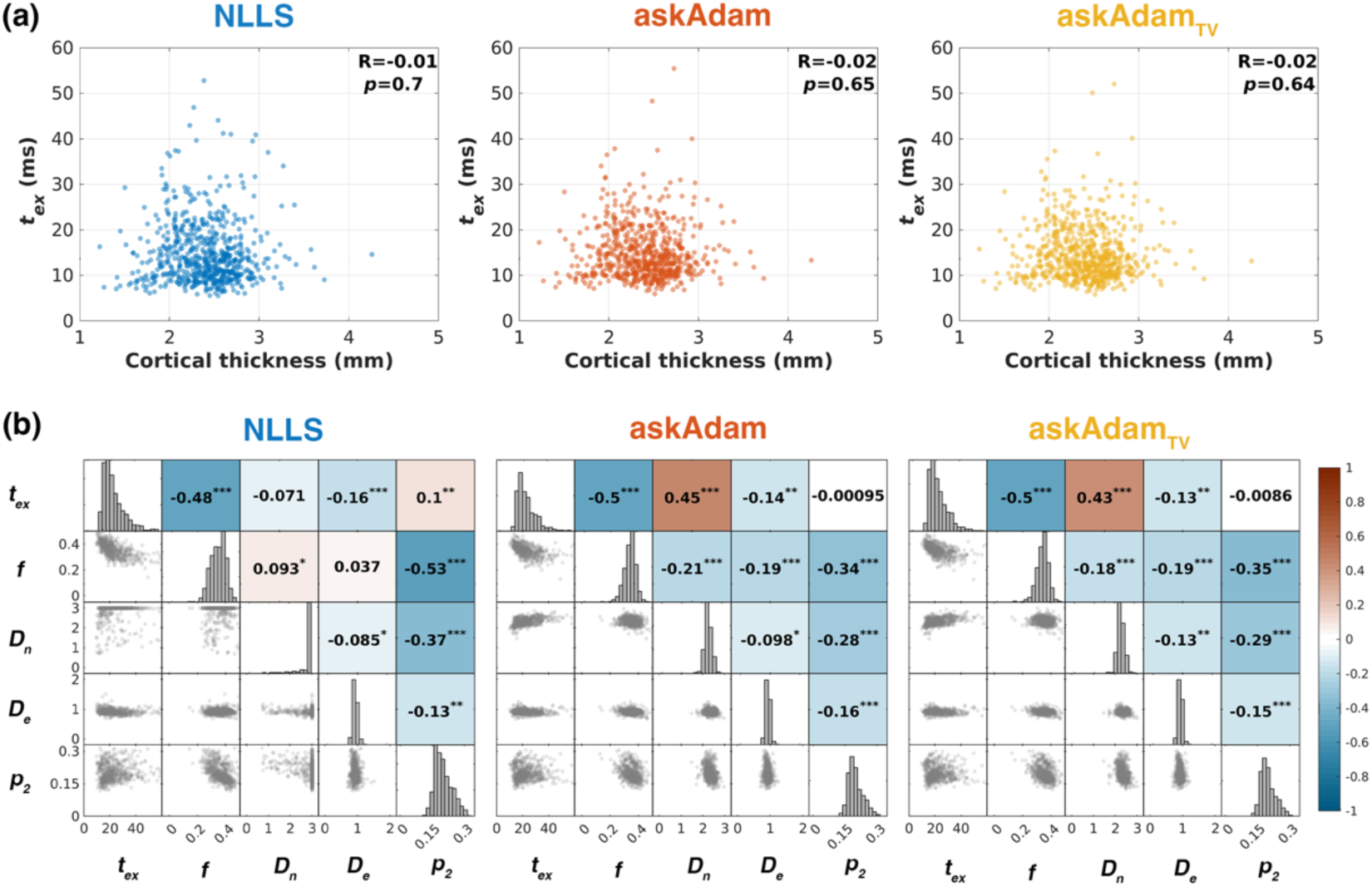
(a) Scatter plots of exchange time *t*_*ex*_ versus cortical thickness across all cortical ROIs and all subjects, with Pearson’s correlation coefficient R and *p*-value are also reported. (b) Correlation matrix between the NEXI microstructure parameters for each solver. \**p*-value<0.05; \*\**p*-value<0.01; \*\*\**p*-value<0.001.

-0.29) were observed for askAdam and askAdam_TV_ (Figure 9b).

## 5. Discussion

In this work, we studied the intercompartmental exchange times between neurites and extracellular water in the living human brain using the NEXI model fitted to data acquired on the state-of-the-art Connectome 2.0 scanner, which enabled us to obtain high-quality, whole-brain dMRI data at strong diffusion weighting (*b*-values up to 17.5 ms/μm^2^) and short diffusion time (*t_min_*=13 ms). Similar to previous animal studies (Jelescu et al., 2022; Olesen et al., 2022), we observed that the zeroth order rationally invariant (spherical mean) dMRI signals decreased with diffusion times at each b-value across the cortex, which was considered as the signature of intercompartmental exchange in grey matter (Olesen et al., 2022). While the measured exchange time across the human cortex using the Connectome 1.0 protocol agreed with the literature with a similar setup (Uhl et al., 2024), the exchange time derived from the Connectome 2.0 protocol was substantially shorter, with median±IQR of approximately 13±8 ms.

The exchange time was observed to vary across the cortical ribbon. Furthermore, we introduced a novel processing tool to accelerate NEXI model fitting by more than 100-fold utilizing the latest GPU computing technology with comparable accuracy and improved precision compared to conventional voxel-wise NLLS fitting.

### Comparison between Connectome 1.0 and Connectome 2.0 protocols

We evaluated the inter-compartmental exchange time in the living human brain at the individual level using the ultra-high-gradient performance Connectome 2.0 scanner. The gradient system of the Connectome 2.0 scanner is five times more powerful than its predecessor Connectome 1.0 (considering together a magnetic field gradient 1.67 times stronger and 3 times faster than Connectome 1.0). This allows the Connectome 2.0 scanner to achieve the shortest diffusion time of 13 ms instead of 21 ms as Connectome 1.0 with similar diffusion weighting, facilitating access to experimental conditions better suited to probing fast water exchange when the two water compartments are less well mixed. Under the ideal conditions (i.e., with Gaussian noise or the noise statistics considered in the signal model), our simulations suggested that both Connectome 1.0 and Connectome 2.0 protocols can perform equally well (Figure 2c). The advantages of the shorter diffusion times accessible on Connectome 2.0 are evident when the exchange time is relatively short (≤20 ms). Compared to the Connectome 1.0 protocol, the estimation of exchange time ≤20 ms using the Connectome 2.0 protocol was more precise and accurate, especially when the noise floor was not accounted for in the signal model (left panel, Figure 2c). This observation is supported by the in vivo exchange time comparison between the two protocols (Figure 5), where the Connectome 1.0 derived exchange time is 2.67 times longer than that of the Connectome 2.0 using the ordinary NEXI model, driven by the reduced estimations in both *f* and *r*_*n*_, whereas the Connectome 1.0 estimations using the NEXI with Rician mean model became substantially shorter and more comparable to the Connectome 2.0 protocol results, potentially due to the incomplete removal of the noise floor by the denoising method. Note that our Connectome 1.0 *in vivo* exchange times derived from the ordinary NEXI model were comparable to the results reported in another study using the same method (103.9 ms using the ordinary NEXI model and 42.3 ms using the NEXI with Rician mean model) (Uhl et al., 2024). Besides the noise floor effect, another hypothesis that may account for the observed differences in exchange times measured on Connectome 2.0 versus Connectome 1.0 could be that water exchange occurs across a broad range of times, as suggested in a recent report (Cai et al., 2023). As a result, using shorter diffusion times may sensitize the measurement of intercompartmental water exchange to the fast exchange process. Previously, shorter exchange times of 3-10 ms were observed in the *ex vivo* rat brain using the SMEX model in an experiment that used relatively short diffusion times (7.5-16 ms) (Olesen et al., 2022). We acknowledge that it is challenging to compare our results directly with those in the *ex vivo* rat brain due to potential interspecies differences in membrane permeability, tissue constituents and water transport mechanisms caused by experiment conditions such as fixation and tissue preparation (Williamson et al., 2023). Differences related to the dMRI acquisition protocol and hardware, including the diffusion times, diffusion weightings and SNR, can also introduce additional variations observed between the two studies. On the other hand, Veraart et al. estimated exchange times of 10-30 ms in the *in vivo* human brain gray matter using the anisotropic Kärger model (Veraart et al., 2020), and Williamson et al. measured an exchange time of 10 ms in the living mouse spinal cord via two-dimensional diffusion exchange spectroscopy (Williamson et al., 2023, 2019), which are both in agreement with our Connectome 2.0 measurements. Other imaging factors, such as SNR, may also affect the measured values. As shown in the noise propagation analysis, the (residual) Rician noise floor can lead to an overestimation of the exchange time (Figure 2c). Even for Gaussian noise, low SNR conditions can lead to slightly overestimated exchange times as indicated by the *in silico* head phantom simulations (Figure 3). Therefore, extra care may be needed on data processing and result interpretation when performing NEXI on standard clinical scanners as the gradient pulse duration, minimum achievable diffusion time and SNR will not be comparable to high-gradient performance systems like Connectome 1.0 or 2.0.

### Implications on noise propagation analysis

Our noise propagation results indicate that the narrow pulse approximation utilized in the NEXI model adequately captures the exchange time given the clinically relevant SNR with high gradient performance setups. This observation remains consistent across different gradient pulse durations (6-10 ms). Although the SMEX model provides a more precise representation of the exchange dMRI signals, its direct usage on NLLS fitting could be relatively challenging due to the longer computational time required for solving the associated ordinary differential equations, especially for whole brain data. Accelerating SMEX forward signal generation using techniques such as artificial neural networks may be valuable to further improve the estimation performance in exchange time measurement in the future. One limitation of our simulation is that we did not take the restricted diffusion from the soma compartment into account for the forward signal simulation, which is also affected by the change of diffusion encoding gradient duration. On the other hand, the Rician noise floor correction has a more notable effect on the accuracy and precision of exchange time estimation. Incorporating the Rician noise model into the NEXI signal model or having Gaussian noise present in the data minimizes estimation bias, aligning with findings from prior research (Uhl et al., 2024). Estimating noise levels in the data is crucial for implementing the NEXI model of Rician mean, achievable through methods such as MP-PCA noise level estimation (Veraart et al., 2016) or through the temporal statistics on multiple *b*=0 images (assuming the noise level is constant over time). Alternatively, techniques like Rician-bias-corrected MP-PCA denoising (Tournier et al., 2023) or using real-value dMRI data (Eichner et al., 2015; Fan et al., 2020; Tian et al., 2022) can reduce the Rician noise floor and potentially achieve noise statistics closer to Gaussian noise conditions. However, it is of the utmost importance to ensure the noise floor is eliminated by using these approaches, as the estimated exchange time is highly sensitive to the residual noise floor, as illustrated in Figure 2c and Figure 5. In our extended analysis, results suggest that higher-order rotational invariants of dMRI signals may not contribute significantly to NEXI microstructure parameter estimation (Figure 3), likely due to the highly disperse nature of neurite orientations in gray matter. In most cases, the zeroth order signals (spherical mean) are sufficient for estimating the exchange time. Nevertheless, incorporating the second-order signals introduces an additional microstructure parameter (*p*_2_), representing neurite orientation dispersion, without sacrificing estimation performance (see also Supplementary Figure S4 and Table S1 for *in vivo* comparison), which could be relevant in disease contexts.

### Effects of diffusion weighting on estimated exchange time

We utilized strong diffusion weighting (maximum *b*-values of 17.5 ms/μm^2^) for NEXI to ensure water exchange can be sampled when the hindered diffusion of extracellular water does not contribute to the diffusion signal. To gain further insight into whether such strong diffusion weighting is needed to measure the exchange time accurately, we performed a retrospective analysis by repeating the ROI-averaged NEXI estimation using dMRI data limited to *b*=6.5 ms/μm^2^. While the exchange times remain comparable in frontal, parietal and occipital gray matter regions, reductions in exchange time are evident in the temporal gray matter, amygdala, and hippocampus when high *b*-value data are excluded (refer to Supplementary Figure S7 and Table S2). This decrease in *t*_*ex*_ = (1 − *f*)/*r*_*n*_ is attributed to the apparent overestimation of the intra-neurite volume fraction *f* (in contrast to using the full b-value range), with the estimated exchange rates *r*_*n*_from neurites to the extracellular space remaining consistent across both datasets.

### *In vivo* exchange time across cortical ribbon

Interestingly, the measured exchange times exhibit spatial variability across the cortex, showing a pattern akin to cortical myelination (Edwards et al., 2018; Marques et al., 2017) derived from multimodality images, as well as to the previous *in vivo* NEXI study (Uhl et al., 2024). Specifically, regions with higher levels of myelination, such as the motor cortex, somatosensory cortex, visual cortex, transverse temporal gyrus, and posterior part of the cingulate cortex, tend to display longer exchange times. Additionally, there appears to be a spatial gradient of myelination increasing from the anterior to the posterior parts of the cingulate cortex, consistent with the observed pattern in the exchange time projection illustrated in Figure 8. Further investigation is needed to elucidate the association between cortical myelination and intercompartmental exchange times.

### Accelerating NEXI processing

We have also demonstrated a significant acceleration of NEXI model fitting, achieving over 100-fold increase in speed using the GPU-based askAdam framework while maintaining comparable estimation performance. Our evaluation using the *in silico* head phantom revealed that, although askAdam’s estimation accuracy is slightly reduced compared to voxel-wise NLLS fitting, the bias in exchange time introduced by askAdam is small (difference on median=0.41 ms at SNR=50 between NLLS and askAdam, Figure 4), particularly when considering inter-subject variability (Figure 8). Another advantage of askAdam is its ability to integrate spatial regularization by updating all microstructure parameters simultaneously during the fitting process. This feature proves particularly beneficial when dealing with low SNR data. As a proof-of-concept demonstration, we implemented 2D-TV regularization on the neurite volume fraction parameter map (see Supplementary Figure S2). Yet, more advanced methods, such as total generalized variation (TGV) (Bredies et al., 2010; Knoll et al., 2011) or structure-guided TV (Ehrhardt and Betcke, 2016), can also be employed. These methods have demonstrated better preservation of anatomical structures, which could be important for cortical gray matter given its highly folded nature. Moreover, the flexibility of askAdam’s loss function allows spatial regularization to extend beyond a single parameter map. It can be applied simultaneously across multi-parameter maps along with other anatomical information, such as sharp edges in the image (Liu et al., 2011). Despite spatial regularization may be useful in reducing the spatial variance and stabilizing the estimation, caution should be exercised to determine the optimal regularization parameter to prevent measurement bias resulting from over-regularization (over-smoothing) that could potentially decrease the sensitivity of the method in assessing specific pathologies, as diseased tissues often have greater microstructural variations when compared to normal tissues. Nonetheless, spatial regularization is only an optional feature of askAdam and can be disabled in the optimization. As shown in Figure 8, the estimation performance without TV regularization is nearly identical to the results obtained with TV regularization at the group-level comparison. It is worth noting that the primary benefit of utilizing the ‘askAdam’ framework lies in its computation time efficiency. Since the estimation principle is still based on minimizing the signal differences between a tissue model and the measurement data, it will inherit similar degeneracy issues in ill-conditioned parameter estimation as in NLLS voxel-wise approaches.

### Limitations

Our study has some limitations. First, the NEXI model employed in our analysis does not explicitly account for water diffusion within the soma. It is established that the soma contributes approximately 10-20% of the water signal at low *b*-values in gray matter (Olesen et al., 2022). At weak diffusion weighting, the signal contribution from the soma may be non-negligible. We observed markedly shorter measured exchange times when incorporating *b*=1 ms/μm^2^ in the NEXI fitting (Supplementary Table S2). The fit residuals also exhibited stronger systematic differences between the NEXI forward signal and the measurements (see Supplementary Figure S6), prompting us to include data with only *b*>1 ms/μm^2^ for the NEXI fitting. Note that the *in vivo* data demonstrated only weak time-dependence at *b*=1 ms/μm^2^ (see Supplementary Figures S5 and S6). This suggests that at the low diffusion weightings, it will be more challenging to disentangle the exchange effect from diffusion. Expanding the NEXI model to encompass a non-exchanging restricted compartment to represent the soma could potentially alleviate signal discrepancies at low *b*-values (Olesen et al., 2022), yet this would introduce additional model parameters that may compromise estimation precision. On the other hand, signal originating from myelinated fibers in grey matter that is not accounted for in the NEXI model, though comprising only a small portion (Shapson-Coe et al., 2024), may become more pronounced at very high *b*-values (e.g., ≥17 ms/μm^2^ for *in vivo*) and introduce biases in the NEXI estimation (Olesen et al., 2022). An *ex vivo* mouse study at 16.4T by Olesen et al. demonstrated that the differences in exchange time with and without considering the myelinated fiber were between 0.3 ms and 0.9 ms with the measured exchange times between 3.9 ms and 5.8 ms. Whether these differences hold for *in vivo* imaging will need further investigation. However, it will be particularly challenging to separate the myelinated fiber signal from the unmyelinated neurite signal in grey matter *in vivo* human experiments, considering the SNR and *b*-values demands. Other signal sources, including microscopic kurtosis (Henriques et al., 2021), effects of the localization regime (Moutal and Grebenkov, 2020), effects of deviation from idealized geometries (Ianus et al., 2021) and effects of dendritic spines (Palombo et al., 2017) among others, that were not accounted for in the NEXI signal model may also contribute to the exchange time biases we observed between the Connectome 1.0 and Connectome 2.0 protocols. While intra-vascular and extra-vascular signal was considered to be highly suppressed due to the use of high *b*-value data here, the exchange effect between these two compartments may also need to be considered when low *b*-value (<1 ms**/**µm^2^) data is used to fit the model with soma compartment.

Another limitation of NEXI is that the signal model does not account for the potential transverse relaxation time differences between the two compartments. Several reports have suggested that the intra-neurite transverse relaxation time is slightly longer than that of the extra-cellular water (Gong et al., 2024; Lee et al., 2024; Veraart et al., 2018). This difference can lead to an echo time dependence on the estimated neurite volume fraction, thus introducing an additional study bias in the estimated exchange time when different echo times are used in the acquisition. This effect may be alleviated by incorporating the compartmental transverse relaxation rates in the NEXI model with dMRI data acquired at multiple echo times (Lee et al., 2024). The additional compartmental relaxation parameters may also provide extra insight into the microstructure environment, but this will certainly lengthen the scan time of the existing protocol due to additional data acquisition.

The drawback of including only higher *b*-value (>1 ms/μm^2^) data is the reduced sensitivity to estimate the neurite diffusivity. Previous works demonstrated that the neurite diffusivity was about 2 μm^2^/ms in white matter (Coelho et al., 2022; Dhital et al., 2019) and gray matter (Ianuş et al., 2022; Palombo et al., 2020), which required lower *b*-value data for a reliable estimation. This is in line with our simulation results and *in vivo* data where the neurite diffusivity is the least robust estimated parameter in the NEXI model given the current imaging protocols and often fitted to the upper bound of the allowed range. Setting a higher fitting upper bound for neurite diffusivity only shows minor impacts on the estimated exchange time (see Supplementary Figure S3). Further investigation is necessary to accurately assess soma properties in conjunction with the intercompartmental exchange effect.

The partial volume effect arising from the superficial white matter and CSF due to the relatively large voxel size compared to cortical thickness can also affect the measurement accuracy (our dMRI data: 2 mm isotropic vs typical cortical thickness=2.2-2.9 mm across all ages (Frangou et al., 2022)). We examined the potential impact of the partial volume effect in our data through correlation analysis between the exchange time and cortical thickness and found no significant association between the two parameters (Figure 9a). While this evidence may suggest that the partial volume effect due to local white matter and CSF may not influence the measured exchange time significantly, it is also possible that they could have an opposite effect in the NEXI estimation. Further investigation is required to understand how different tissue types (grey matter, white matter and CSF) coexist in a voxel may impact the estimation of the *in vivo* NEXI exchange time. Increasing the spatial resolution can mitigate the partial volume effect at the expense of reducing SNR and lengthening scan time. Nevertheless, given the high-quality data provided by the Connectome 2.0 scanner (SNR=55 at *b*=0 before denoising), obtaining high-resolution dMRI while maintaining reasonable SNR using standard spin-echo EPI sequences may be feasible. Alternatively, recent advancements in pulse sequence development (Dong et al., 2024; Ramos-Llordén et al., 2020; Setsompop et al., 2018), denoising (Henriques et al., 2023; Lemberskiy et al., 2023; Olesen et al., 2023), super-resolution techniques (Coupé et al., 2013; H. H. Lee et al., 2019; Manjón et al., 2010) and diffusion sampling strategy (Fan et al., 2017) for high-resolution diffusion imaging offer a promising avenue to achieve high resolution with good SNR for gray matter imaging. Another advantage of higher spatial resolution is the potential to investigate the intercompartment exchange effect at multiple cortical depths, providing more specific information reflecting tissue composition and arrangement across the cortical layers.

To be able to obtain high-quality data for NEXI exchange time mapping, we employed a protocol comprising 15 *b*-values at 64 diffusion gradient directions from 3 diffusion times, resulting in more than 80 min total acquisition time, which is clearly too long for clinical applications. Reducing the number of diffusion gradient directions by half as the protocol used for Connectome 1.0 and Connectome 2.0 comparison can shorten the scan time to 40 min while retaining comparable results as the 64-direction high SNR protocol. Future studies will focus on protocol optimization on sequence parameters, such as diffusion times, *b*-values, and number of diffusion gradient directions, to achieve a more accessible scan time (<20 min) for clinical and research applications.

## 6. Conclusions

The ultra-high-gradient performance Connectome 2.0 scanner provides high-quality data and greater flexibility in experimental design to measure intercompartmental exchange effects between intracellular and extracellular water in the brain gray matter. The *in vivo* intercompartmental exchange time was estimated to be about 13±8 ms across the human cortex based on the anisotropic Kärger model. Our work further indicated that addressing the Rician noise floor sufficiently, either in the signal model or in the data processing, is crucial to measure the short exchange time (≤20 ms) using NEXI, especially when a short diffusion time cannot be easily deployed due to hardware constraints. The narrow pulse approximation in NEXI did not introduce significant bias in the exchange time measurements using (ultra-)high gradient performance MRI systems, such as Connectome 1.0 and 2.0 scanners, under clinically relevant SNR circumstances. Using the GPU-based askAdam processing pipeline, we accelerated the NEXI data fitting process by 100-fold. Our work provides additional insights and groundwork that may aid in NEXI protocol optimization for *in vivo* imaging in future studies.

## Data and Code Availability

The data and code that support the findings of this study will be made available upon reasonable request. The GPU processing tool for the ordinary NEXI model and NEXI with Rician mean model parameter estimation is available at https://github.com/kschan0214/gacelle.

## Author Contributions

**Kwok-Shing Chan**: Conceptualization, Methodology, Software, Validation, Formal analysis, Investigation, Writing, Visualization

**Yixin Ma**: Investigation, Writing

**Hansol Lee**: Investigation, Writing

**José P. Marques**: Software, Conceptualization, Writing

**Jonas Olesen**: Software, Conceptualization, Methodology

**Santiago Coelho**: Conceptualization, Methodology, Writing

**Dmitry S Novikov**: Conceptualization, Methodology, Writing

**Sune Jespersen**: Software, Conceptualization, Methodology, Writing

**Susie Y. Huang**: Conceptualization, Methodology, Writing, Supervision, Funding acquisition

**Hong-Hsi Lee**: Conceptualization, Methodology, Software, Writing, Supervision, Funding acquisition

## Declaration of Competing Interests

The authors have no competing interests related to the findings of this work.

### Acknowledgements

This study was support by the Office of the Director (OD) of the National Institutes of Health (NIH) in partnership with the National Institute of Dental and Craniofacial Research (NIDCR) under award number DP5OD031854, the OD of the NIH under award number S10OD032184, the National Institute of Neurological Disorders and Stroke (NINDS) of NIH under award numbers R01NS118187 and R01NS088040, the National Institute of Biomedical Imaging and Bioengineering (NIBIB) under award numbers P41EB015896, P41EB030006, U01EB026996 and P41EB017183, the National Center for Research Resources (NCRR) or NIH under award numbers S10RR023401 and S10RR019307, and the National Institute on Aging under award number R21AG085795. SJ acknowledges financial support from the Lundbeck Foundation (R396-2022-183). SJ is also supported by the Danish National Research Foundation (CFIN), and the Danish Ministry of Science, Innovation, and Education (MINDLab).

## References

Ades-Aron, B., Veraart, J., Kochunov, P., McGuire, S., Sherman, P., Kellner, E., Novikov, D.S., Fieremans, E., 2018. Evaluation of the accuracy and precision of the diffusion parameter EStImation with Gibbs and NoisE removal pipeline. NeuroImage 183, 532–543. 10.1016/j.neuroimage.2018.07.066

Andersson, J.L.R., Skare, S., Ashburner, J., 2003. How to correct susceptibility distortions in spin-echo echo-planar images: application to diffusion tensor imaging. Neuroimage 20, 870–888. 10.1016/s1053-8119(03)00336-7

Andersson, J.L.R., Sotiropoulos, S.N., 2016. An integrated approach to correction for off-resonance effects and subject movement in diffusion MR imaging. Neuroimage 125, 1063–1078. 10.1016/j.neuroimage.2015.10.019

Åslund, I., Nowacka, A., Nilsson, M., Topgaard, D., 2009. Filter-exchange PGSE NMR determination of cell membrane permeability. J. Magn. Reson. 200, 291–295. 10.1016/j.jmr.2009.07.015

Assaf, Y., Blumenfeld-Katzir, T., Yovel, Y., Basser, P.J., 2008. Axcaliber: A method for measuring axon diameter distribution from diffusion MRI. Magn. Reson. Med. 59, 1347–1354. 10.1002/mrm.21577

Assaf, Y., Freidlin, R.Z., Rohde, G.K., Basser, P.J., 2004. New modeling and experimental framework to characterize hindered and restricted water diffusion in brain white matter. Magn. Reson. Med. 52, 965–978. 10.1002/mrm.20274

Avants, B.B., Tustison, N.J., Song, G., Cook, P.A., Klein, A., Gee, J.C., 2011. A reproducible evaluation of ANTs similarity metric performance in brain image registration. Neuroimage 54, 2033–2044. 10.1016/j.neuroimage.2010.09.025

Badaut, J., Fukuda, A.M., Jullienne, A., Petry, K.G., 2014. Aquaporin and brain diseases. Biochim. Biophys. Acta (BBA) - Gen. Subj. 1840, 1554–1565. 10.1016/j.bbagen.2013.10.032

Bai, R., Li, Z., Sun, C., Hsu, Y.-C., Liang, H., Basser, P., 2020. Feasibility of filter-exchange imaging (FEXI) in measuring different exchange processes in human brain. NeuroImage 219, 117039. 10.1016/j.neuroimage.2020.117039

Bai, R., Springer, C.S., Plenz, D., Basser, P.J., 2019. Brain active transmembrane water cycling measured by MR is associated with neuronal activity. Magn. Reson. Med. 81, 1280–1295. 10.1002/mrm.27473

Bai, R., Springer, C.S., Plenz, D., Basser, P.J., 2018. Fast, Na+/K+ pump driven, steady-state transcytolemmal water exchange in neuronal tissue: A study of rat brain cortical cultures. Magn. Reson. Med. 79, 3207–3217. 10.1002/mrm.26980

Basser, P.J., 1997. New Histological and Physiological Stains Derived from Diffusion-Tensor MR Images. Ann. N. York Acad. Sci. 820, 123–138. 10.1111/j.1749-6632.1997.tb46192.x

Beaulieu, C., 2002. The basis of anisotropic water diffusion in the nervous system - a technical review. NMR in biomedicine 15, 435–455. 10.1002/nbm.782

Billot, B., Magdamo, C., Cheng, Y., Arnold, S.E., Das, S., Iglesias, J.E., 2023. Robust machine learning segmentation for large-scale analysis of heterogeneous clinical brain MRI datasets. Proc National Acad Sci 120, e2216399120. 10.1073/pnas.2216399120

Bredies, K., Kunisch, K., Pock, T., 2010. Total Generalized Variation. SIAM J. Imaging Sci. 3, 492–526. 10.1137/090769521

Cai, T.X., Benjamini, D., Komlosh, M.E., Basser, P.J., Williamson, N.H., 2018. Rapid detection of the presence of diffusion exchange. J. Magn. Reson. 297, 17–22. 10.1016/j.jmr.2018.10.004

Cai, T.X., Williamson, N.H., Ravin, R., Basser, P.J., 2023. Multiexponential analysis of diffusion exchange times reveals a distinct exchange process associated with metabolic activity, in: Proceedings 31. Annual Meeting International Society for Magnetic Resonance in Medicine. Toronto, Canada, p. 5017.

Callaghan, P.T., Furó, I., 2004. Diffusion-diffusion correlation and exchange as a signature for local order and dynamics. J. Chem. Phys. 120, 4032–4038. 10.1063/1.1642604

Carr, H.Y., Purcell, E.M., 1954. Effects of Diffusion on Free Precession in Nuclear Magnetic Resonance Experiments. Phys. Rev. 94, 630–638. 10.1103/physrev.94.630

Chakwizira, A., Zhu, A., Foo, T., Westin, C.-F., Szczepankiewicz, F., Nilsson, M., 2023. Diffusion MRI with free gradient waveforms on a high-performance gradient system: Probing restriction and exchange in the human brain. NeuroImage 283, 120409. 10.1016/j.neuroimage.2023.120409

Chan, K.-S., Kim, T.H., Bilgic, B., Marques, J.P., 2022. Semi-supervised learning for fast multi-compartment relaxometry myelin water imaging (MCR-MWI), in: Proceedings 30. Annual Meeting International Society for Magnetic Resonance in Medicine. London, United Kingdom, p. 1639. 10.58530/2022/1639

Coelho, S., Baete, S.H., Lemberskiy, G., Ades-Aron, B., Barrol, G., Veraart, J., Novikov, D.S., Fieremans, E., 2022. Reproducibility of the Standard Model of diffusion in white matter on clinical MRI systems. NeuroImage 257, 119290. 10.1016/j.neuroimage.2022.119290

Coupé, P., Manjón, J.V., Chamberland, M., Descoteaux, M., Hiba, B., 2013. Collaborative patch-based super-resolution for diffusion-weighted images. NeuroImage 83, 245–261. 10.1016/j.neuroimage.2013.06.030

Dhital, B., Reisert, M., Kellner, E., Kiselev, V.G., 2019. Intra-axonal diffusivity in brain white matter. NeuroImage 189, 543–550. 10.1016/j.neuroimage.2019.01.015

Dong, Z., Reese, T.G., Lee, H.-H., Huang, S.Y., Polimeni, J.R., Wald, L.L., Wang, F., 2024. Romer-EPTI: Rotating-view motion-robust super-resolution EPTI for SNR-efficient distortion-free in-vivo mesoscale dMRI and microstructure imaging. bioRxiv 2024.01.26.577343. 10.1101/2024.01.26.577343

Edwards, L.J., Kirilina, E., Mohammadi, S., Weiskopf, N., 2018. Microstructural imaging of human neocortex in vivo. Neuroimage 182, 184–206. 10.1016/j.neuroimage.2018.02.055

Ehrhardt, M.J., Betcke, M.M., 2016. Multicontrast MRI Reconstruction with Structure-Guided Total Variation. SIAM J. Imaging Sci. 9, 1084–1106. 10.1137/15m1047325

Eichner, C., Cauley, S.F., Cohen-Adad, J., Möller, H.E., Turner, R., Setsompop, K., Wald, L.L., 2015. Real diffusion-weighted MRI enabling true signal averaging and increased diffusion contrast. NeuroImage 122, 373–384. 10.1016/j.neuroimage.2015.07.074

Fan, Q., Nummenmaa, A., Polimeni, J.R., Witzel, T., Huang, S.Y., Wedeen, V.J., Rosen, B.R., Wald, L.L., 2017. HIgh b-value and high Resolution Integrated Diffusion (HIBRID) imaging. NeuroImage 150, 162–176. 10.1016/j.neuroimage.2017.02.002

Fan, Q., Nummenmaa, A., Witzel, T., Ohringer, N., Tian, Q., Setsompop, K., Klawiter, E.C., Rosen, B.R., Wald, L.L., Huang, S.Y., 2020. Axon diameter index estimation independent of fiber orientation distribution using high-gradient diffusion MRI. NeuroImage 222, 117197. 10.1016/j.neuroimage.2020.117197

Fan, Q., Witzel, T., Nummenmaa, A., Dijk, K.R.A.V., Horn, J.D.V., Drews, M.K., Somerville, L.H., Sheridan, M.A., Santillana, R.M., Snyder, J., Hedden, T., Shaw, E.E., Hollinshead, M.O., Renvall, V., Zanzonico, R., Keil, B., Cauley, S., Polimeni, J.R., Tisdall, D., Buckner, R.L., Wedeen, V.J., Wald, L.L., Toga, A.W., Rosen, B.R., 2016. MGH–USC Human Connectome Project datasets with ultra-high b-value diffusion MRI. NeuroImage 124, 1108–1114. 10.1016/j.neuroimage.2015.08.075

Feinberg, D.A., Beckett, A.J.S., Vu, A.T., Stockmann, J., Huber, L., Ma, S., Ahn, S., Setsompop, K., Cao, X., Park, S., Liu, C., Wald, L.L., Polimeni, J.R., Mareyam, A., Gruber, B., Stirnberg, R., Liao, C., Yacoub, E., Davids, M., Bell, P., Rummert, E., Koehler, M., Potthast, A., Gonzalez-Insua, I., Stocker, S., Gunamony, S., Dietz, P., 2023. Next-generation MRI scanner designed for ultra-high-resolution human brain imaging at 7 Tesla. Nat. Methods 20, 2048–2057. 10.1038/s41592-023-02068-7

Fieremans, E., Jensen, J.H., Helpern, J.A., 2011. White matter characterization with diffusional kurtosis imaging. Neuroimage 58, 177–188. 10.1016/j.neuroimage.2011.06.006

Fieremans, E., Novikov, D.S., Jensen, J.H., Helpern, J.A., 2010. Monte Carlo study of a two-compartment exchange model of diffusion. NMR Biomed. 23, 711–724. 10.1002/nbm.1577

Fischl, B., 2012. FreeSurfer. NeuroImage 62, 774–781. 10.1016/j.neuroimage.2012.01.021

Foo, T.K.F., Tan, E.T., Vermilyea, M.E., Hua, Y., Fiveland, E.W., Piel, J.E., Park, K., Ricci, J., Thompson, P.S., Graziani, D., Conte, G., Kagan, A., Bai, Y., Vasil, C., Tarasek, M., Yeo, D.T.B., Snell, F., Lee, D., Dean, A., DeMarco, J.K., Shih, R.Y., Hood, M.N., Chae, H., Ho, V.B., 2020. Highly efficient head-only magnetic field insert gradient coil for achieving simultaneous high gradient amplitude and slew rate at 3.0T (MAGNUS) for brain microstructure imaging. Magn. Reson. Med. 83, 2356–2369. 10.1002/mrm.28087

Frangou, S., Modabbernia, A., Williams, S.C.R., Papachristou, E., Doucet, G.E., Agartz, I., Aghajani, M., Akudjedu, T.N., Albajes-Eizagirre, A., Alnæs, D., Alpert, K.I., Andersson, M., Andreasen, N.C., Andreassen, O.A., Asherson, P., Banaschewski, T., Bargallo, N., Baumeister, S., Baur-Streubel, R., Bertolino, A., Bonvino, A., Boomsma, D.I., Borgwardt, S., Bourque, J., Brandeis, D., Breier, A., Brodaty, H., Brouwer, R.M., Buitelaar, J.K., Busatto, G.F., Buckner, R.L., Calhoun, V., Canales-Rodríguez, E.J., Cannon, D.M., Caseras, X., Castellanos, F.X., Cervenka, S., Chaim-Avancini, T.M., Ching, C.R.K., Chubar, V., Clark, V.P., Conrod, P., Conzelmann, A., Crespo-Facorro, B., Crivello, F., Crone, E.A., Dale, A.M., Dannlowski, U., Davey, C., Geus, E.J.C. de, Haan, L. de, Zubicaray, G.I. de, Braber, A. den, Dickie, E.W., Giorgio, A.D., Doan, N.T., Dørum, E.S., Ehrlich, S., Erk, S., Espeseth, T., Fatouros-Bergman, H., Fisher, S.E., Fouche, J., Franke, B., Frodl, T., Fuentes-Claramonte, P., Glahn, D.C., Gotlib, I.H., Grabe, H., Grimm, O., Groenewold, N.A., Grotegerd, D., Gruber, O., Gruner, P., Gur, R.E., Gur, R.C., Hahn, T., Harrison, B.J., Hartman, C.A., Hatton, S.N., Heinz, A., Heslenfeld, D.J., Hibar, D.P., Hickie, I.B., Ho, B., Hoekstra, P.J., Hohmann, S., Holmes, A.J., Hoogman, M., Hosten, N., Howells, F.M., Pol, H.E.H., Huyser, C., Jahanshad, N., James, A., Jernigan, T.L., Jiang, J., Jönsson, E.G., Joska, J.A., Kahn, R., Kalnin, A., Kanai, R., Klein, M., Klyushnik, T.P., Koenders, L., Koops, S., Krämer, B., Kuntsi, J., Lagopoulos, J., Lázaro, L., Lebedeva, I., Lee, W.H., Lesch, K., Lochner, C., Machielsen, M.W.J., Maingault, S., Martin, N.G., Martínez-Zalacaín, I., Mataix-Cols, D., Mazoyer, B., McDonald, C., McDonald, B.C., McIntosh, A.M., McMahon, K.L., McPhilemy, G., Meinert, S., Menchón, J.M., Medland, S.E., Meyer-Lindenberg, A., Naaijen, J., Najt, P., Nakao, T., Nordvik, J.E., Nyberg, L., Oosterlaan, J., Foz, V.O. de la, Paloyelis, Y., Pauli, P., Pergola, G., Pomarol-Clotet, E., Portella, M.J., Potkin, S.G., Radua, J., Reif, A., Rinker, D.A., Roffman, J.L., Rosa, P.G.P., Sacchet, M.D., Sachdev, P.S., Salvador, R., Sánchez-Juan, P., Sarró, S., Satterthwaite, T.D., Saykin, A.J., Serpa, M.H., Schmaal, L., Schnell, K., Schumann, G., Sim, K., Smoller, J.W., Sommer, I., Soriano-Mas, C., Stein, D.J., Strike, L.T., Swagerman, S.C., Tamnes, C.K., Temmingh, H.S., Thomopoulos, S.I., Tomyshev, A.S., Tordesillas-Gutiérrez, D., Trollor, J.N., Turner, J.A., Uhlmann, A., Heuvel, O.A. van den, Meer, D. van den, Wee, N.J.A. van der, Haren, N.E.M. van, Ent, D. van ’t, Erp, T.G.M. van, Veer, I.M., Veltman, D.J., Voineskos, A., Völzke, H., Walter, H., Walton, E., Wang, L., Wang, Y., Wassink, T.H., Weber, B., Wen, W., West, J.D., Westlye, L.T., Whalley, H., Wierenga, L.M., Wittfeld, K., Wolf, D.H., Worker, A., Wright, M.J., Yang, K., Yoncheva, Y., Zanetti, M.V., Ziegler, G.C., (KaSP), K.S.P., Thompson, P.M., Dima, D., 2022. Cortical thickness across the lifespan: Data from 17,075 healthy individuals aged 3–90 years. Hum. Brain Mapp. 43, 431–451. 10.1002/hbm.25364

Gong, T., Tax, C.M., Mancini, M., Jones, D.K., Zhang, H., Palombo, M., 2024. Multi-TE SANDI: Quantifying compartmental T2 relaxation times in the grey matter, in: Proceedings 32. Annual Meeting International Society for Magnetic Resonance in Medicine. p. 0766.

Harms, R.L., Fritz, F.J., Tobisch, A., Goebel, R., Roebroeck, A., 2017. Robust and fast nonlinear optimization of diffusion MRI microstructure models. NeuroImage 155, 82–96. 10.1016/j.neuroimage.2017.04.064

Henriques, R.N., Ianuş, A., Novello, L., Jovicich, J., Jespersen, S.N., Shemesh, N., 2023. Efficient PCA denoising of spatially correlated redundant MRI data. Imaging Neurosci. 1, 1–26. 10.1162/imag_a_00049

Henriques, R.N., Jespersen, S.N., Shemesh, N., 2021. Evidence for microscopic kurtosis in neural tissue revealed by correlation tensor MRI. Magn. Reson. Med. 86, 3111–3130. 10.1002/mrm.28938

Hernandez-Fernandez, M., Reguly, I., Jbabdi, S., Giles, M., Smith, S., Sotiropoulos, S.N., 2019. Using GPUs to accelerate computational diffusion MRI: From microstructure estimation to tractography and connectomes. NeuroImage 188, 598–615. 10.1016/j.neuroimage.2018.12.015

Huang, S.Y., Tian, Q., Fan, Q., Witzel, T., Wichtmann, B., McNab, J.A., Bireley, J.D., Machado, N., Klawiter, E.C., Mekkaoui, C., Wald, L.L., Nummenmaa, A., 2020. High-gradient diffusion MRI reveals distinct estimates of axon diameter index within different white matter tracts in the in vivo human brain. Brain Struct. Funct. 225, 1277–1291. 10.1007/s00429-019-01961-2

Huang, S.Y., Witzel, T., Keil, B., Scholz, A., Davids, M., Dietz, P., Rummert, E., Ramb, R., Kirsch, J.E., Yendiki, A., Fan, Q., Tian, Q., Ramos-Llordén, G., Lee, H.-H., Nummenmaa, A., Bilgic, B., Setsompop, K., Wang, F., Avram, A.V., Komlosh, M., Benjamini, D., Magdoom, K.N., Pathak, S., Schneider, W., Novikov, D.S., Fieremans, E., Tounekti, S., Mekkaoui, C., Augustinack, J., Berger, D., Shapson-Coe, A., Lichtman, J., Basser, P.J., Wald, L.L., Rosen, B.R., 2021. Connectome 2.0: Developing the next-generation ultra-high gradient strength human MRI scanner for bridging studies of the micro-, meso- and macro-connectome. NeuroImage 243, 118530. 10.1016/j.neuroimage.2021.118530

Ianus, A., Alexander, D.C., Zhang, H., Palombo, M., 2021. Mapping complex cell morphology in the grey matter with double diffusion encoding MR: A simulation study. Neuroimage 241, 118424. 10.1016/j.neuroimage.2021.118424

Ianuş, A., Carvalho, J., Fernandes, F.F., Cruz, R., Chavarrias, C., Palombo, M., Shemesh, N., 2022. Soma and Neurite Density MRI (SANDI) of the in-vivo mouse brain and comparison with the Allen Brain Atlas. NeuroImage 254, 119135. 10.1016/j.neuroimage.2022.119135

Jelescu, I.O., Skowronski, A. de, Geffroy, F., Palombo, M., Novikov, D.S., 2022. Neurite Exchange Imaging (NEXI): A minimal model of diffusion in gray matter with inter-compartment water exchange. Neuroimage 256, 119277. 10.1016/j.neuroimage.2022.119277

Jespersen, S.N., Kroenke, C.D., Østergaard, L., Ackerman, J.J.H., Yablonskiy, D.A., 2007. Modeling dendrite density from magnetic resonance diffusion measurements. Neuroimage 34, 1473–1486. 10.1016/j.neuroimage.2006.10.037

Jovicich, J., Czanner, S., Greve, D., Haley, E., Kouwe, A. van der, Gollub, R., Kennedy, D., Schmitt, F., Brown, G., MacFall, J., Fischl, B., Dale, A., 2006. Reliability in multi-site structural MRI studies: Effects of gradient non-linearity correction on phantom and human data. NeuroImage 30, 436–443. 10.1016/j.neuroimage.2005.09.046

Jung, S., Lee, H., Ryu, K., Song, J.E., Park, M., Moon, W., Kim, D., 2021. Artificial neural network for multi-echo gradient echo–based myelin water fraction estimation. Magnet Reson Med 85, 380–389. 10.1002/mrm.28407

Kaden, E., Kelm, N.D., Carson, R.P., Does, M.D., Alexander, D.C., 2016. Multi-compartment microscopic diffusion imaging. Neuroimage 139, 346–359. 10.1016/j.neuroimage.2016.06.002

Kärger, J., 1985. NMR self-diffusion studies in heterogeneous systems. Adv. Colloid Interface Sci. 23, 129–148. 10.1016/0001-8686(85)80018-x

Kellner, E., Dhital, B., Kiselev, V.G., Reisert, M., 2016. Gibbs-ringing artifact removal based on local subvoxel-shifts. Magnetic resonance in medicine 76, 1574–1581. 10.1002/mrm.26054

Khateri, M., Reisert, M., Sierra, A., Tohka, J., Kiselev, V.G., 2022. What does FEXI measure? NMR Biomed. 35, e4804. 10.1002/nbm.4804

King, L.S., Kozono, D., Agre, P., 2004. From structure to disease: the evolving tale of aquaporin biology. Nat. Rev. Mol. Cell Biol. 5, 687–698. 10.1038/nrm1469

Kingma, D.P., Ba, J., 2014. Adam: A Method for Stochastic Optimization. arXiv. 10.48550/arxiv.1412.6980

Klein, A., Tourville, J., 2012. 101 Labeled Brain Images and a Consistent Human Cortical Labeling Protocol. Front. Neurosci. 6, 171. 10.3389/fnins.2012.00171

Knoll, F., Bredies, K., Pock, T., Stollberger, R., 2011. Second order total generalized variation (TGV) for MRI. Magn. Reson. Med. 65, 480–491. 10.1002/mrm.22595

Lampinen, B., Lätt, J., Wasselius, J., Westen, D. van, Nilsson, M., 2021. Time dependence in diffusion MRI predicts tissue outcome in ischemic stroke patients. Magn. Reson. Med. 86, 754–764. 10.1002/mrm.28743

Lasič, S., Nilsson, M., Lätt, J., Ståhlberg, F., Topgaard, D., 2011. Apparent exchange rate mapping with diffusion MRI. Magn. Reson. Med. 66, 356–365. 10.1002/mrm.22782

Lätt, J., Nilsson, M., Westen, D. van, Wirestam, R., Ståhlberg, F., Brockstedt, S., 2009. Diffusion-weighted MRI measurements on stroke patients reveal water-exchange mechanisms in sub-acute ischaemic lesions. NMR Biomed. 22, 619– 628. 10.1002/nbm.1376

Le Bihan, D., 1995. Molecular diffusion, tissue microdynamics and microstructure. NMR Biomed. 8, 375–386. 10.1002/nbm.1940080711

Lee, H.-H., Chan, K.S., Llorden, G.R., Huang, S.Y., 2024. TENEXI: Echo time-dependent neurite exchange imaging for in vivo evaluation of exchange time and relaxation time on the Connectome 2.0 scanner, in: Proceedings 32. Annual Meeting International Society for Magnetic Resonance in Medicine. p. 3468.

Lee, H.H., Lin, Y.C., Lemberskiy, G., Ades-Aron, B., Baete, S., Boada, F.E., Fieremans, E., Novikov, D.S., 2019. SUper-REsolution TRACTography (SURE-TRACT) pipeline using self-similarity between diffusional and anatomical images, in: Proceedings 27. Annual Meeting International Society for Magnetic Resonance in Medicine. Montreal, Canada, p. 0167.

Lee, H.H., Olesen, J.L., Tian, Q., Llorden, G.R., Jespersen, S.N., Huang, S.Y., 2022. Revealing diffusion time-dependence and exchange effect in the in vivo human brain gray matter by using high gradient diffusion MRI, in: Proceedings 30. Annual Meeting International Society for Magnetic Resonance in Medicine. London, United Kingdom, p. 0254. 10.58530/2022/0254

Lee, H.-H., Papaioannou, A., Novikov, D.S., Fieremans, E., 2020. In vivo observation and biophysical interpretation of time-dependent diffusion in human cortical gray matter. Neuroimage 222, 117054. 10.1016/j.neuroimage.2020.117054

Lee, J., Lee, D., Choi, J.Y., Shin, D., Shin, H., Lee, Jongho, 2019. Artificial neural network for myelin water imaging. Magn. Reson. Med. 83, 1875–1883. 10.1002/mrm.28038

Lemberskiy, G., Chandarana, H., Bruno, M., Ginocchio, L.A., Huang, C., Tong, A., Keerthivasan, M.B., Fieremans, E., Novikov, D.S., 2023. Feasibility of Accelerated Prostate Diffusion-Weighted Imaging on 0.55 T MRI Enabled With Random Matrix Theory Denoising. Investig. Radiol. 58, 720–729. 10.1097/rli.0000000000000979

Li, C., Fieremans, E., Novikov, D.S., Ge, Y., Zhang, J., 2023. Measuring water exchange on a preclinical MRI system using filter exchange and diffusion time dependent kurtosis imaging. Magn. Reson. Med. 89, 1441–1455. 10.1002/mrm.29536

Liu, T., Liu, J., Rochefort, L. de, Spincemaille, P., Khalidov, I., Ledoux, J.R., Wang, Y., 2011. Morphology enabled dipole inversion (MEDI) from a single-angle acquisition: Comparison with COSMOS in human brain imaging. Magnetic resonance in medicine 66, 777–783. 10.1002/mrm.22816

Ma, D., Gulani, V., Seiberlich, N., Liu, K., Sunshine, J.L., Duerk, J.L., Griswold, M.A., 2013. Magnetic resonance fingerprinting. Nature 495, 187–192. 10.1038/nature11971

MacAulay, N., 2021. Molecular mechanisms of brain water transport. Nat. Rev. Neurosci. 22, 326–344. 10.1038/s41583-021-00454-8

Mahmutovic, M., Shrestha, M., Ramos-Llordén, G., Scholz, A., Kirsch, J.E., Wald, L.L., Möller, H.E., Mekkaoui, C., Huang, S.Y., Keil, B., 2024. A 72-channel Head Coil with an Integrated 16-Channel Field Camera for the Connectome 2.0 Scanner, in: Proceedings 32. Annual Meeting International Society for Magnetic Resonance in Medicine. Singapore, p. 1030.

Manjón, J.V., Coupé, P., Buades, A., Collins, D.L., Robles, M., 2010. MRI Superresolution Using Self-Similarity and Image Priors. Int. J. Biomed. Imaging 2010, 425891. 10.1155/2010/425891

Marques, J.P., Hollander, D.D., Norris, D.G., Chan, K.-S., 2023. Improved R2* and QSM mapping for dummies - ask Adam, in: Proceedings 31. Annual Meeting International Society for Magnetic Resonance in Medicine. Toronto, Canada, p. 1089.

Marques, J.P., Khabipova, D., Gruetter, R., 2017. Studying cyto and myeloarchitecture of the human cortex at ultra-high field with quantitative imaging: R1, R2 * and magnetic susceptibility. NeuroImage 147, 152–163. 10.1016/j.neuroimage.2016.12.009

Martins, J.P. de A., Nilsson, M., Lampinen, B., Palombo, M., While, P.T., Westin, C.-F., Szczepankiewicz, F., 2021. Neural networks for parameter estimation in microstructural MRI: Application to a diffusion-relaxation model of white matter. NeuroImage 244, 118601. 10.1016/j.neuroimage.2021.118601

Moutal, N., Grebenkov, D.S., 2020. The localization regime in a nutshell. J. Magn. Reson. 320, 106836. 10.1016/j.jmr.2020.106836

Nedjati-Gilani, G.L., Schneider, T., Hall, M.G., Cawley, N., Hill, I., Ciccarelli, O., Drobnjak, I., Wheeler-Kingshott, C.A.M.G., Alexander, D.C., 2017. Machine learning based compartment models with permeability for white matter microstructure imaging. NeuroImage 150, 119–135. 10.1016/j.neuroimage.2017.02.013

Nilsson, M., Lätt, J., Westen, D. van, Brockstedt, S., Lasič, S., Ståhlberg, F., Topgaard, D., 2013. Noninvasive mapping of water diffusional exchange in the human brain using filter-exchange imaging. Magn. Reson. Med. 69, 1572–1580. 10.1002/mrm.24395

Ning, L., Nilsson, M., Lasič, S., Westin, C.-F., Rathi, Y., 2018. Cumulant expansions for measuring water exchange using diffusion MRI. J. Chem. Phys. 148, 074109. 10.1063/1.5014044

Novikov, D.S., Fieremans, E., Jespersen, S.N., Kiselev, V.G., 2019. Quantifying brain microstructure with diffusion MRI: Theory and parameter estimation. NMR Biomed. 32, e3998. 10.1002/nbm.3998

Novikov, D.S., Kiselev, V.G., Jespersen, S.N., 2018a. On modeling. Magnet Reson Med 79, 3172–3193. 10.1002/mrm.27101

Novikov, D.S., Veraart, J., Jelescu, I.O., Fieremans, E., 2018b. Rotationally-invariant mapping of scalar and orientational metrics of neuronal microstructure with diffusion MRI. NeuroImage 174, 518–538. 10.1016/j.neuroimage.2018.03.006

Olesen, J.L., Ianus, A., Østergaard, L., Shemesh, N., Jespersen, S.N., 2023. Tensor denoising of multidimensional MRI data. Magn. Reson. Med. 89, 1160–1172. 10.1002/mrm.29478

Olesen, J.L., Østergaard, L., Shemesh, N., Jespersen, S.N., 2022. Diffusion time dependence, power-law scaling, and exchange in gray matter. Neuroimage 251, 118976. 10.1016/j.neuroimage.2022.118976

Orton, M.R., Collins, D.J., Koh, D., Leach, M.O., 2014. Improved intravoxel incoherent motion analysis of diffusion weighted imaging by data driven Bayesian modeling. Magn. Reson. Med. 71, 411–420. 10.1002/mrm.24649

Palombo, M., Ianus, A., Guerreri, M., Nunes, D., Alexander, D.C., Shemesh, N., Zhang, H., 2020. SANDI: A compartment-based model for non-invasive apparent soma and neurite imaging by diffusion MRI. Neuroimage 215, 116835. 10.1016/j.neuroimage.2020.116835

Palombo, M., Ligneul, C., Hernandez-Garzon, E., Valette, J., 2017. Can we detect the effect of spines and leaflets on the diffusion of brain intracellular metabolites? Neuroimage 182, 283–293. 10.1016/j.neuroimage.2017.05.003

Papadopoulos, M.C., Verkman, A.S., 2013. Aquaporin water channels in the nervous system. Nat. Rev. Neurosci. 14, 265–277. 10.1038/nrn3468

Pasternak, O., Assaf, Y., Intrator, N., Sochen, N., 2008. Variational multiple-tensor fitting of fiber-ambiguous diffusion-weighted magnetic resonance imaging voxels. Magn. Reson. Imaging 26, 1133–1144. 10.1016/j.mri.2008.01.006

QSM Consensus Organization Committee, Bilgic, B., Costagli, M., Chan, K., Duyn, J., Langkammer, C., Lee, J., Li, X., Liu, C., Marques, J.P., Milovic, C., Robinson, S.D., Schweser, F., Shmueli, K., Spincemaille, P., Straub, S., Zijl, P. van, Wang, Y., ISMRM Electro-Magnetic Tissue Properties Study Group, 2024. Recommended implementation of quantitative susceptibility mapping for clinical research in the brain: A consensus of the ISMRM electro-magnetic tissue properties study group. Magn. Reson. Med. 91, 1834–1862. 10.1002/mrm.30006

Ramos-Llordén, G., Ning, L., Liao, C., Mukhometzianov, R., Michailovich, O., Setsompop, K., Rathi, Y., 2020. High-fidelity, accelerated whole-brain submillimeter in vivo diffusion MRI using gSlider-spherical ridgelets (gSlider-SR). Magn. Reson. Med. 84, 1781–1795. 10.1002/mrm.28232

Reisert, M., Kellner, E., Dhital, B., Hennig, J., Kiselev, V.G., 2017. Disentangling micro from mesostructure by diffusion MRI: A Bayesian approach. NeuroImage 147, 964–975. 10.1016/j.neuroimage.2016.09.058

Setsompop, K., Fan, Q., Stockmann, J., Bilgic, B., Huang, S., Cauley, S.F., Nummenmaa, A., Wang, F., Rathi, Y., Witzel, T., Wald, L.L., 2018. High-resolution in vivo diffusion imaging of the human brain with generalized slice dithered enhanced resolution: Simultaneous multislice (gSlider-SMS). Magn. Reson. Med. 79, 141–151. 10.1002/mrm.26653

Setsompop, K., Kimmlingen, R., Eberlein, E., Witzel, T., Cohen-Adad, J., McNab, J.A., Keil, B., Tisdall, M.D., Hoecht, P., Dietz, P., Cauley, S.F., Tountcheva, V., Matschl, V., Lenz, V.H., Heberlein, K., Potthast, A., Thein, H., Horn, J.V., Toga, A., Schmitt, F., Lehne, D., Rosen, B.R., Wedeen, V., Wald, L.L., 2013. Pushing the limits of in vivo diffusion MRI for the Human Connectome Project. Neuroimage 80, 220–233. 10.1016/j.neuroimage.2013.05.078

Shapson-Coe, A., Januszewski, M., Berger, D.R., Pope, A., Wu, Y., Blakely, T., Schalek, R.L., Li, P.H., Wang, S., Maitin-Shepard, J., Karlupia, N., Dorkenwald, S., Sjostedt, E., Leavitt, L., Lee, D., Troidl, J., Collman, F., Bailey, L., Fitzmaurice, A., Kar, R., Field, B., Wu, H., Wagner-Carena, J., Aley, D., Lau, J., Lin, Z., Wei, D., Pfister, H., Peleg, A., Jain, V., Lichtman, J.W., 2024. A petavoxel fragment of human cerebral cortex reconstructed at nanoscale resolution. Science 384, eadk4858. 10.1126/science.adk4858

Springer, C.S., Baker, E.M., Li, X., Moloney, B., Pike, M.M., Wilson, G.J., Anderson, V.C., Sammi, M.K., Garzotto, M.G., Kopp, R.P., Coakley, F.V., Rooney, W.D., Maki, J.H., 2023a. Metabolic activity diffusion imaging (MADI): II. Noninvasive, high-resolution human brain mapping of sodium pump flux and cell metrics. NMR Biomed. 36, e4782. 10.1002/nbm.4782

Springer, C.S., Baker, E.M., Li, X., Moloney, B., Wilson, G.J., Pike, M.M., Barbara, T.M., Rooney, W.D., Maki, J.H., 2023b. Metabolic activity diffusion imaging (MADI): I. Metabolic, cytometric modeling and simulations. NMR Biomed. 36, e4781. 10.1002/nbm.4781

Stejskal, E.O., Tanner, J.E., 1965. Spin Diffusion Measurements: Spin Echoes in the Presence of a Time-Dependent Field Gradient. J Chem Phys 42, 288–292. 10.1063/1.1695690

Tian, Q., Fan, Q., Witzel, T., Polackal, M.N., Ohringer, N.A., Ngamsombat, C., Russo, A.W., Machado, N., Brewer, K., Wang, F., Setsompop, K., Polimeni, J.R., Keil, B., Wald, L.L., Rosen, B.R., Klawiter, E.C., Nummenmaa, A., Huang, S.Y., 2022. Comprehensive diffusion MRI dataset for in vivo human brain microstructure mapping using 300 mT/m gradients. Sci. Data 9, 7. 10.1038/s41597-021-01092-6

Tournier, J.-D., Jeurissen, B., Christiaens, D., 2023. Iterative model-based Rician bias correction and its application to denoising in diffusion MRI, in: Proceedings 31. Annual Meeting International Society for Magnetic Resonance in Medicine. Toronto, Canada, p. 3795.

Tournier, J.-D., Smith, R., Raffelt, D., Tabbara, R., Dhollander, T., Pietsch, M., Christiaens, D., Jeurissen, B., Yeh, C.-H., Connelly, A., 2019. MRtrix3: A fast, flexible and open software framework for medical image processing and visualisation. Neuroimage 202, 116137. 10.1016/j.neuroimage.2019.116137

Uhl, Q., Pavan, T., Molendowska, M., Jones, D.K., Palombo, M., Jelescu, I.O., 2024. Quantifying human gray matter microstructure using neurite exchange imaging (NEXI) and 300 mT/m gradients. Imaging Neurosci. 2, 1–19. 10.1162/imag_a_00104

Veraart, J., Fieremans, E., Novikov, D.S., 2019. On the scaling behavior of water diffusion in human brain white matter. NeuroImage 185, 379–387. 10.1016/j.neuroimage.2018.09.075

Veraart, J., Novikov, D.S., Christiaens, D., Ades-aron, B., Sijbers, J., Fieremans, E., 2016. Denoising of diffusion MRI using random matrix theory. Neuroimage 142, 394–406. 10.1016/j.neuroimage.2016.08.016

Veraart, J., Novikov, D.S., Fieremans, E., 2018. TE dependent Diffusion Imaging (TEdDI) distinguishes between compartmental T 2 relaxation times. NeuroImage 182, 360–369. 10.1016/j.neuroimage.2017.09.030

Veraart, J., Nunes, D., Rudrapatna, U., Fieremans, E., Jones, D.K., Novikov, D.S., Shemesh, N., 2020. Nonivasive quantification of axon radii using diffusion MRI. Elife 9, e49855. 10.7554/elife.49855

Weiger, M., Overweg, J., Rösler, M.B., Froidevaux, R., Hennel, F., Wilm, B.J., Penn, A., Sturzenegger, U., Schuth, W., Mathlener, M., Borgo, M., Börnert, P., Leussler, C., Luechinger, R., Dietrich, B.E., Reber, J., Brunner, D.O., Schmid, T., Vionnet, L., Pruessmann, K.P., 2018. A high-performance gradient insert for rapid and short-T2 imaging at full duty cycle. Magn. Reson. Med. 79, 3256–3266. 10.1002/mrm.26954

Williamson, N.H., Ravin, R., Benjamini, D., Merkle, H., Falgairolle, M., O’Donovan, M.J., Blivis, D., Ide, D., Cai, T.X., Ghorashi, N.S., Bai, R., Basser, P.J., 2019. Magnetic resonance measurements of cellular and sub-cellular membrane structures in live and fixed neural tissue. eLife 8, e51101. 10.7554/elife.51101

Williamson, N.H., Ravin, R., Cai, T.X., Benjamini, D., Falgairolle, M., O’Donovan, M.J., Basser, P.J., 2020. Real-time measurement of diffusion exchange rate in biological tissue. J. Magn. Reson. 317, 106782. 10.1016/j.jmr.2020.106782

Williamson, N.H., Ravin, R., Cai, T.X., Falgairolle, M., O’Donovan, M.J., Basser, P.J., 2023. Water exchange rates measure active transport and homeostasis in neural tissue. PNAS Nexus 2, pgad056. 10.1093/pnasnexus/pgad056

Zhang, H., Schneider, T., Wheeler-Kingshott, C.A., Alexander, D.C., 2012. NODDI: practical in vivo neurite orientation dispersion and density imaging of the human brain. Neuroimage 61, 1000–1016. 10.1016/j.neuroimage.2012.03.072

Zimmerman, J.R., Brittin, W.E., 1957. Nuclear Magnetic Resonance Studies in Multiple Phase Systems: Lifetime of a Water Molecule in an Adsorbing Phase on Silica Gel. J. Phys. Chem. 61, 1328–1333. 10.1021/j150556a015

