## Supplementary for "*In vivo* human neurite exchange imaging (NEXI) at 500 mT/m diffusion gradients"

**Supplementary Materials**

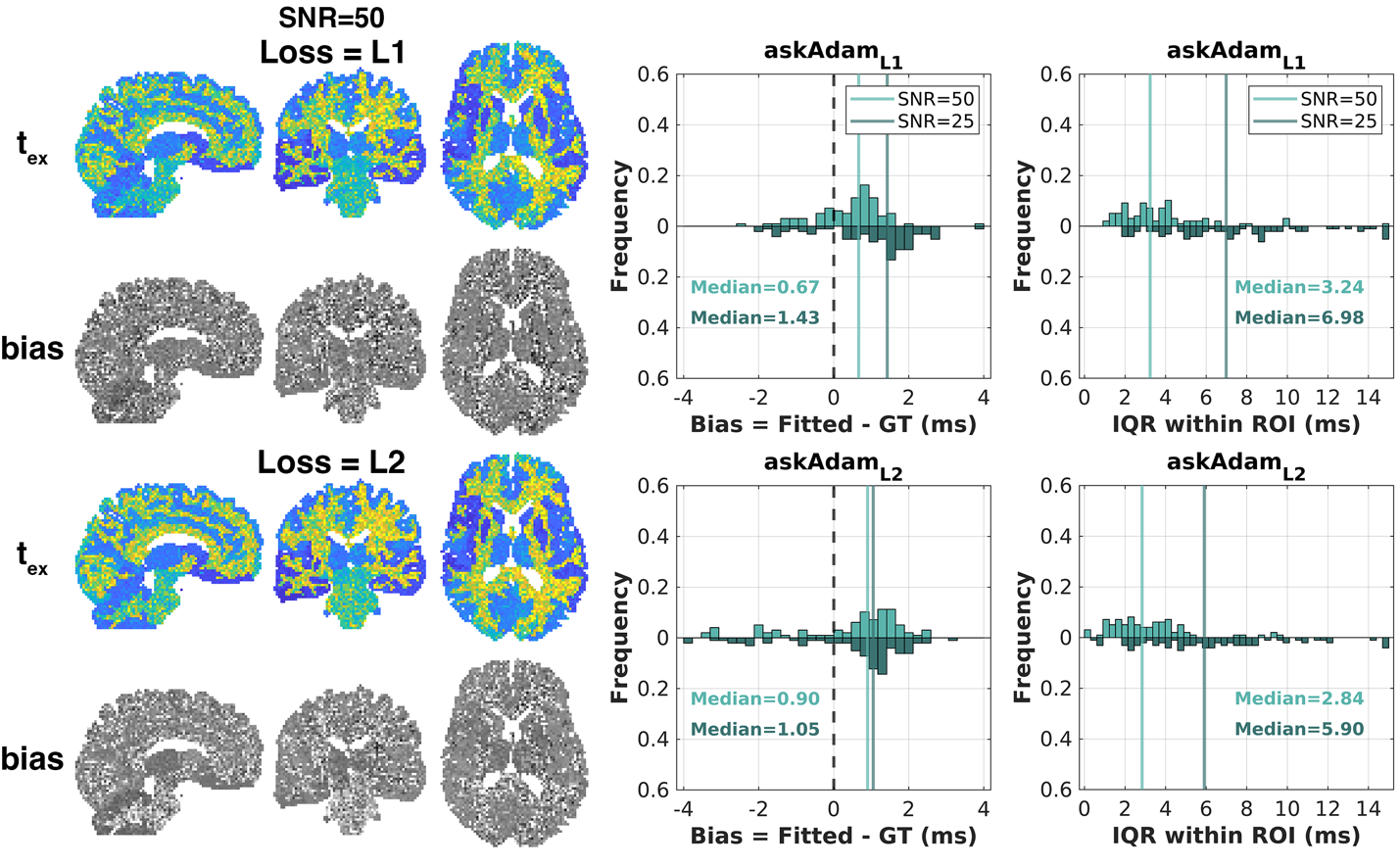

Figure S1: The same result shown in Figure 3 using askAdam solver with (top) L1-norm and (bottom) L2=norm as the loss function as provided in Eq. (7). The L1-norm loss function produces more accurate results at SNR=50, whereas the results of L2-norm are more accurate at SNR=25. The $t_{ex}$ estimation precision is slightly better with the L2-norm. Given our in vivo data have SNR>50 at *b*=0, we decided to use the L1-norm as the loss function in this work.

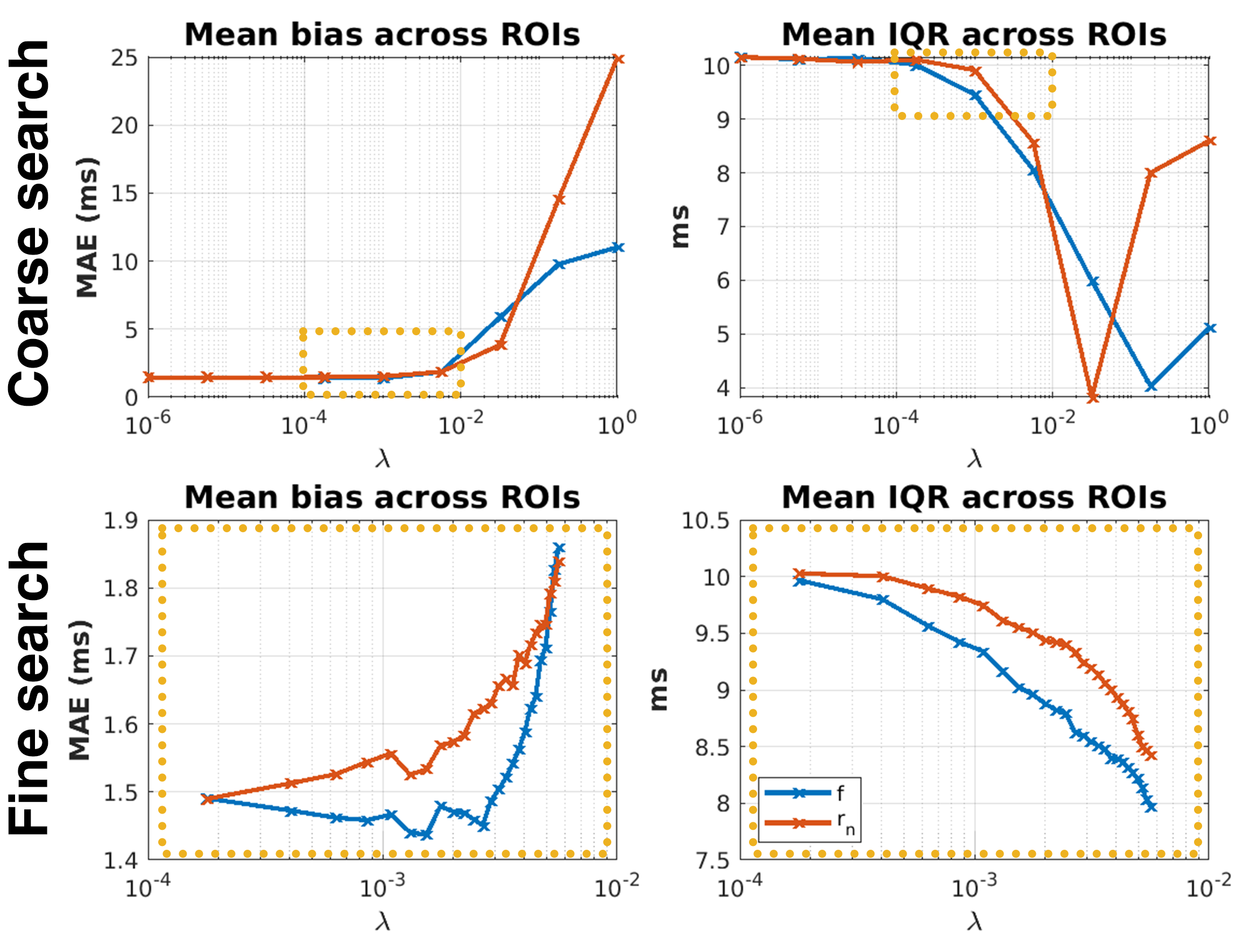

Figure S2: Determining the optimal regularization parameter and map for askAdam_TV_ using *in silico* head phantom data at SNR=50. Regularization on two NEXI parameter maps ($f$ and $r_{n}$) was studied over a range of regularization values. The optimal regularization values were determined in a two-step searching approach: firstly, a coarse search was performed in the range between 1e-6 and 1 with a non-linear step size to sample 9 different regularization values (top row). Mean absolute error (MAE) and the mean IQR of the exchange time across all ROIs were used to decide a narrower range (yellow box) for a finer search. The coarse search results suggested that the regularization is in a good balance of estimation accuracy and precision in the range between 1.8e-4 and 0.056 for both parameters. In the fine-search step, 25 regularization values were sampled over this range with a linear step size of 2.3e-4 (bottom row). Across this narrow range when the estimation accuracy has not deteriorated too severely, applying regularization on the $f$ map is most always performed better than $r_{n}$ (smaller bias and narrower IQR).

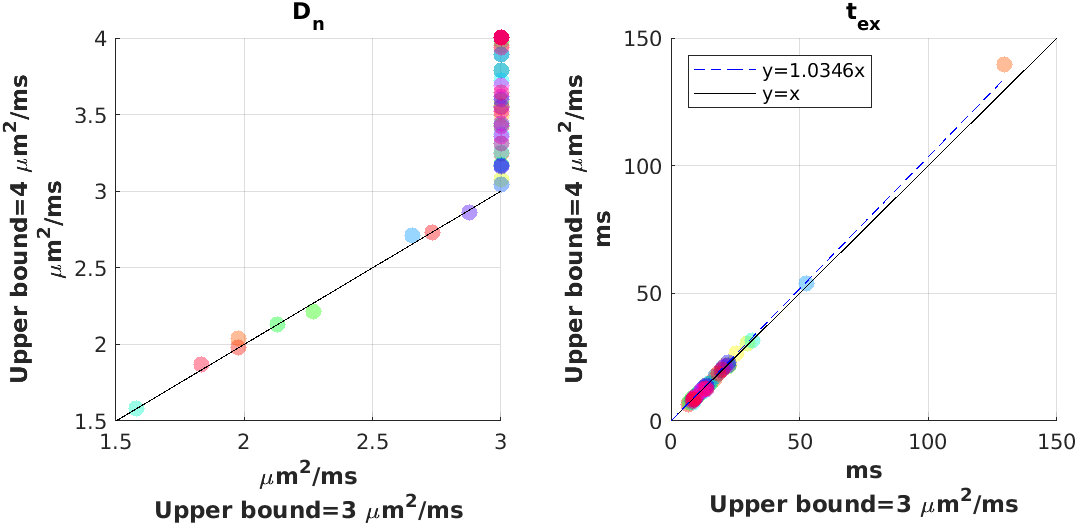

Figure S3: Scatterplots of (left) neurite diffusivity $D_{n}$ and (right) exchange time $t_{ex}$ by setting fitting upper bounds of neurite diffusivity at 3 and 4 μm^2^/ms. Each data point represents the median value of an ROI.

We repeated the *in vivo* ROI fitting analysis described in Section 3.2.3 using zeroth and second-order rotationally invariant dMRI signals, respectively, on the Connectome 2.0 data to validate the aforementioned noise propagation analysis finding. Both $l_{max}=0$ and $l_{max}=2$ NEXI models showed similar fitting quality (dashed lines, Figure S3) on the zeroth order rotationally invariant $S_{l=0}$signal. In this analysis, the exchange time $t_{ex}$ ranged from 14.9 ms (frontal) to 33.5 ms (occipital) for the NEXI $l_{max}=0$ model (Table S1). The values of the fitted parameters were generally similar between $l_{max}=0$ and $l_{max}=2$. In all ROIs, the intra-neurite diffusivity $D_{n}$ was faster than the extra-cellular diffusivity $D_{e}$ for either $l_{max}=0$ or $l_{max}=2$ model fitting.

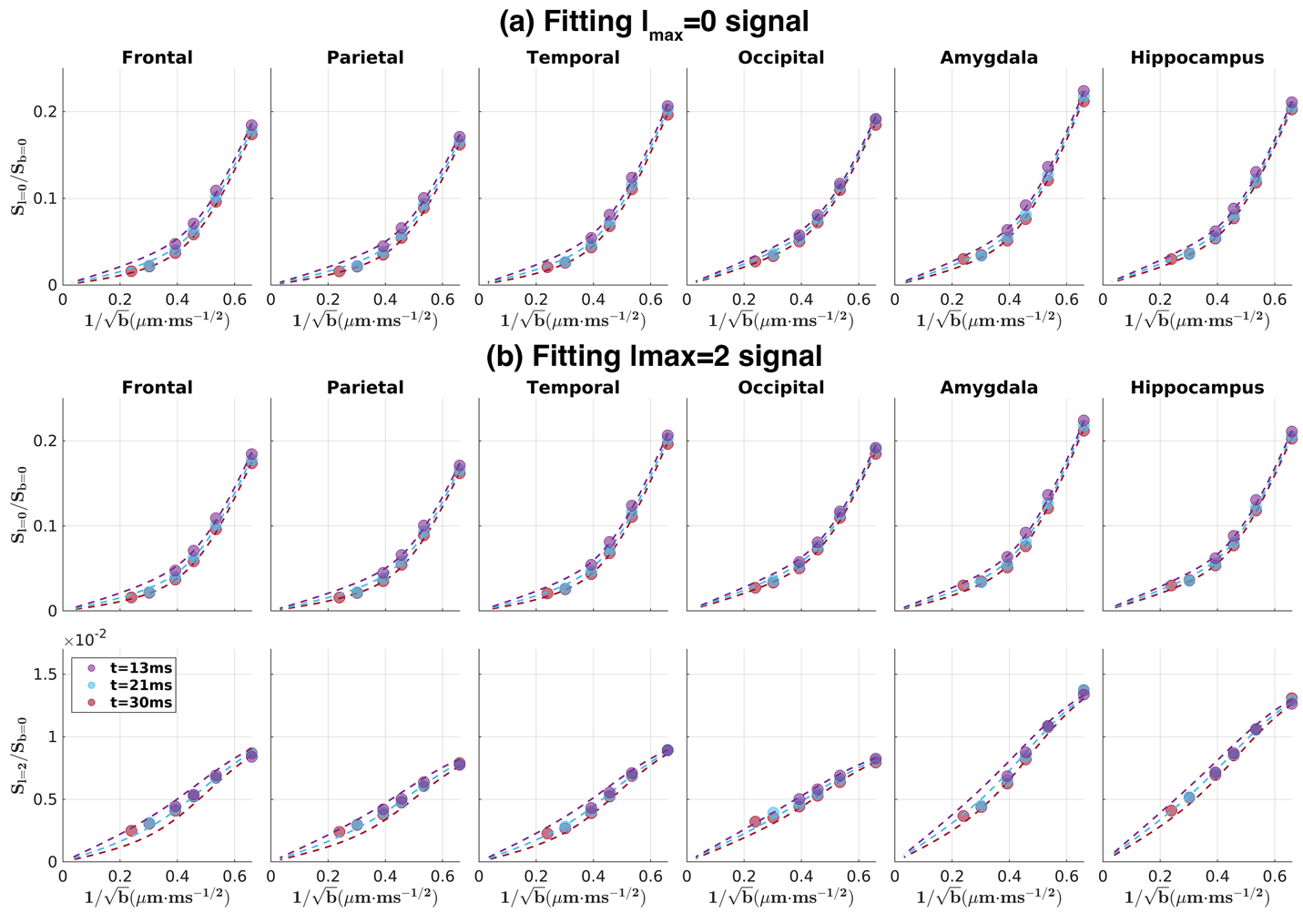

Figure S4: Results of NEXI model fitting on ROI-averaged *in vivo* dMRI signals using Connectome 2.0 protocol data. The NEXI model of $l_{max}=0$ was fitted to (a) only the zeroth order rotationally invariant signals $S_{l=0}$, and the NEXI model of $l_{max}=2$ was fitted to (b) both zeroth and second order rotationally invariant signals, $S_{l=0}$ and $S_{l=2}$. Circles represent the measured data and dashed lines represent the NEXI model fitting curve derived from the fitted tissue parameters.

Table S1: NEXI tissue parameters of grey matter ROI derived from ROI-averaged *in vivo* dMRI signals using Connectome 2.0 data.

|  | $t_{ex}$ (ms) | | $f$ | | $D_{n}$ (𝜇m^2^/ms) | | $D_{e}$ (𝜇m^2^/ms) | | $p_{2}$ |
| --- | --- | --- | --- | --- | --- | --- | --- | --- | --- |
|  | $l_{max}=0$ | $l_{max}=2$ | $l_{max}=0$ | $l_{max}=2$ | $l_{max}=0$ | $l_{max}=2$ | $l_{max}=0$ | $l_{max}=2$ | $l_{max}=2$ |
| Frontal | 14.88 | 14.93 | 0.35 | 0.35 | 3.00 | 3.00 | 0.89 | 0.89 | 0.21 |
| Parietal | 17.79 | 17.84 | 0.32 | 0.32 | 3.00 | 3.00 | 0.94 | 0.94 | 0.21 |
| Temporal | 17.06 | 17.06 | 0.35 | 0.35 | 3.00 | 3.00 | 0.81 | 0.81 | 0.20 |
| Occipital | 33.45 | 33.51 | 0.34 | 0.34 | 3.00 | 3.00 | 0.89 | 0.89 | 0.19 |
| Amygdala | 23.00 | 22.96 | 0.37 | 0.37 | 3.00 | 3.00 | 0.77 | 0.77 | 0.28 |
| Hippocampus | 29.31 | 29.36 | 0.36 | 0.36 | 3.00 | 3.00 | 0.82 | 0.82 | 0.28 |

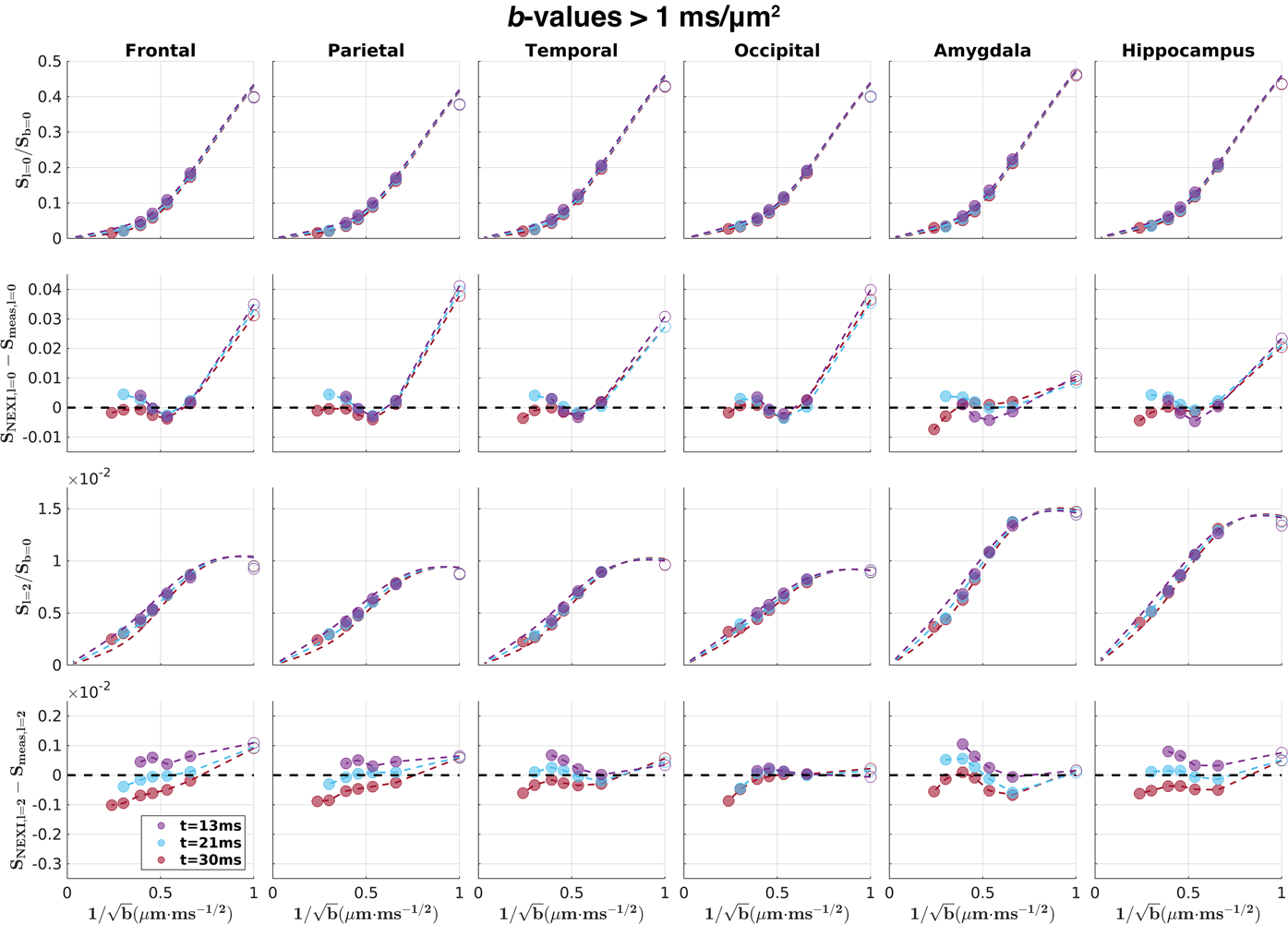

Figure S5: NEXI model fitting with ROI-averaged dMRI signals using $l_{max}=2$ on Connectome 2.0 protocol data. This is the same result as shown in Figure 5a in the main text, but the forward simulated NEXI signal was extrapolated to *b*=1 ms/μm^2^. Solid circles: measured data used in data fitting; hollow circles: measured data that was not used in data fitting. Dashed lines: forward NEXI signals derived from the fitted microstructure parameters.

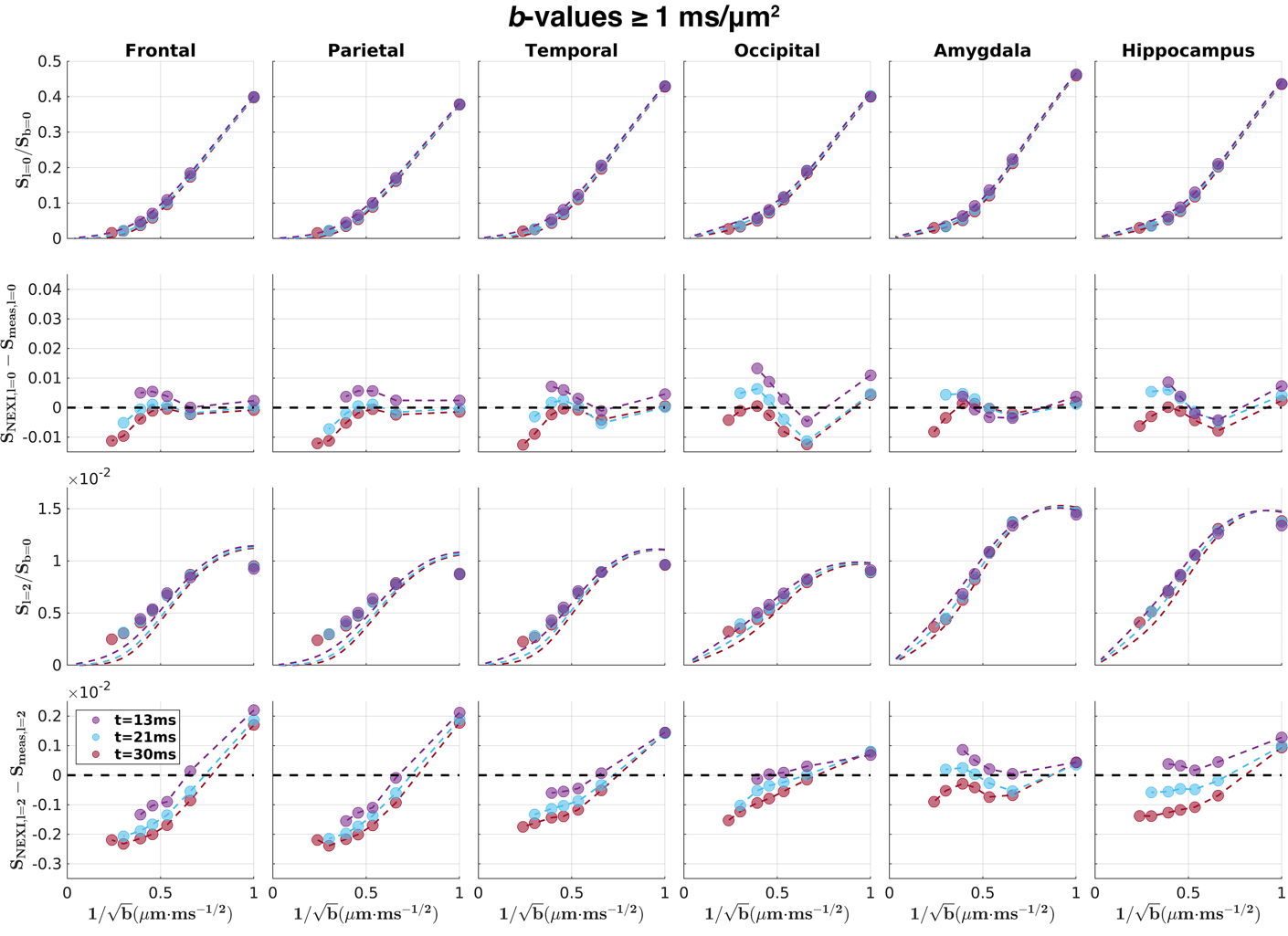

Figure S6: NEXI model fitting with ROI-averaged dMRI signals using $l_{max}=2$ on Connectome 2.0 data. The results shown here were computed with the inclusion of data acquired at *b*=1 ms/μm^2^. Solid circles: measured data used in data fitting. Dashed lines: forward NEXI signals derived from the fitted microstructure parameters.

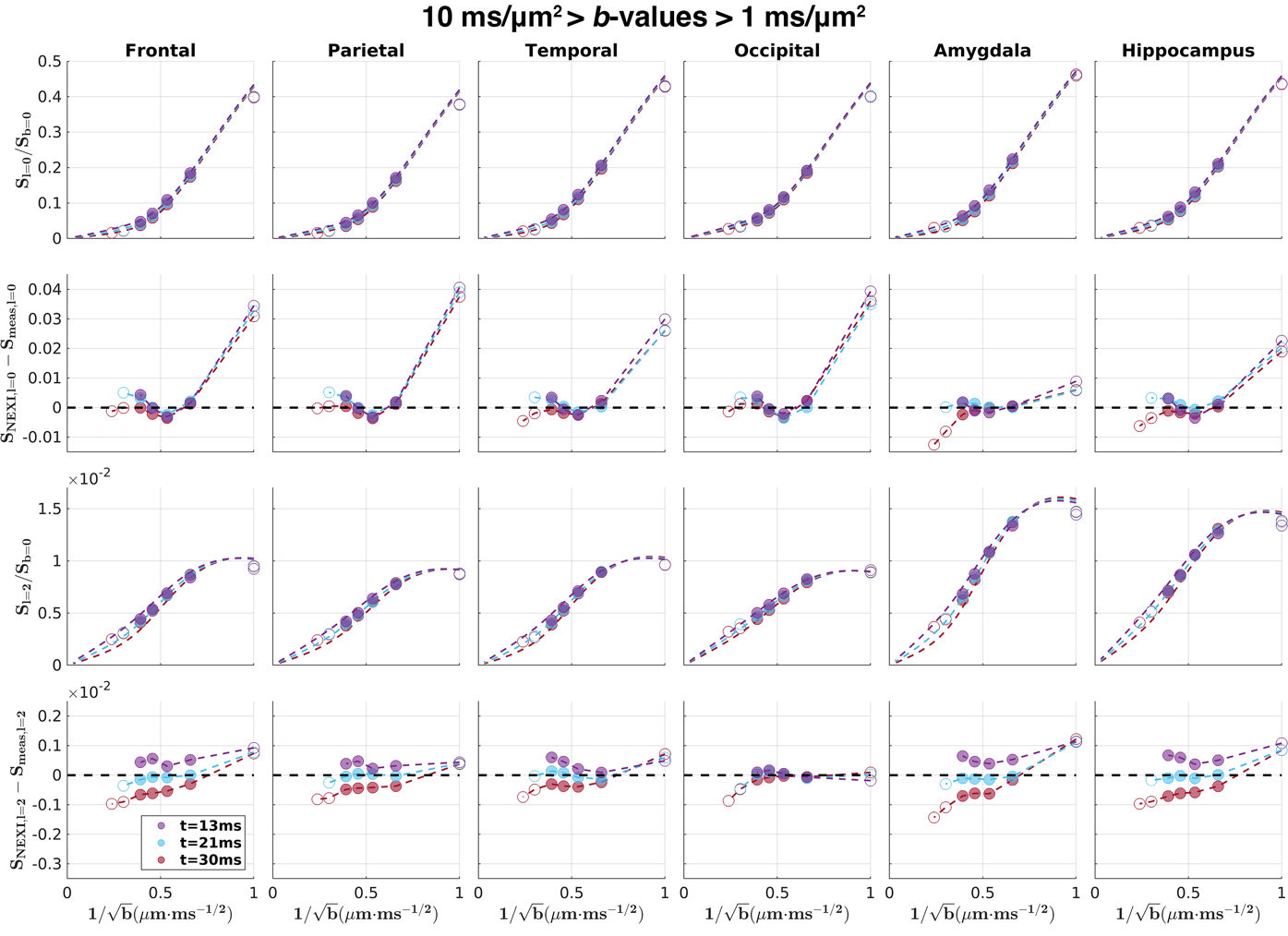

Figure S7: NEXI model fitting with ROI-averaged dMRI signals using $l_{max}=2$ on Connectome 2.0 protocol data. The results shown here were computed when high b-value data were excluded in data fitting (*b_max_*=6.5 ms/μm^2^). Solid circles: measured data used in data fitting; hollow circles: measured data that was not used in data fitting. Dashed lines: forward NEXI signals derived from the fitted microstructure parameters.

Table S2: NEXI microstructure parameters of various grey matter ROIs derived from ROI-averaged dMRI signal using Connectome 2.0 protocol data with different *b*-values.

|  | $t_{ex}$ (ms) | | $f$ | | $D_{n}$ (𝜇m^2^/ms) | | $D_{e}$ (𝜇m^2^/ms) | | $p_{2}$ | |
| --- | --- | --- | --- | --- | --- | --- | --- | --- | --- | --- |
|  | $b\geq1$ | $10>b>1$ | $b\geq1$ | $10>b>1$ | $b\geq1$ | $10>b>1$ | $b\geq1$ | $10>b>1$ | $b\geq1$ | $10>b>1$ |
| Frontal | 1.85 | 15.66 | 0.70 | 0.34 | 3.00 | 3.00 | 1.34 | 0.90 | 0.13 | 0.21 |
| Parietal | 1.13 | 19.35 | 0.78 | 0.31 | 3.00 | 3.00 | 1.89 | 0.94 | 0.12 | 0.21 |
| Temporal | 3.44 | 14.95 | 0.61 | 0.37 | 3.00 | 3.00 | 0.98 | 0.81 | 0.13 | 0.19 |
| Occipital | 14.30 | 34.66 | 0.48 | 0.34 | 3.00 | 3.00 | 1.07 | 0.89 | 0.15 | 0.19 |
| Amygdala | 17.76 | 12.90 | 0.40 | 0.42 | 3.00 | 3.00 | 0.79 | 0.77 | 0.26 | 0.26 |
| Hippocampus | 16.49 | 23.00 | 0.45 | 0.38 | 3.00 | 3.00 | 0.90 | 0.82 | 0.24 | 0.27 |

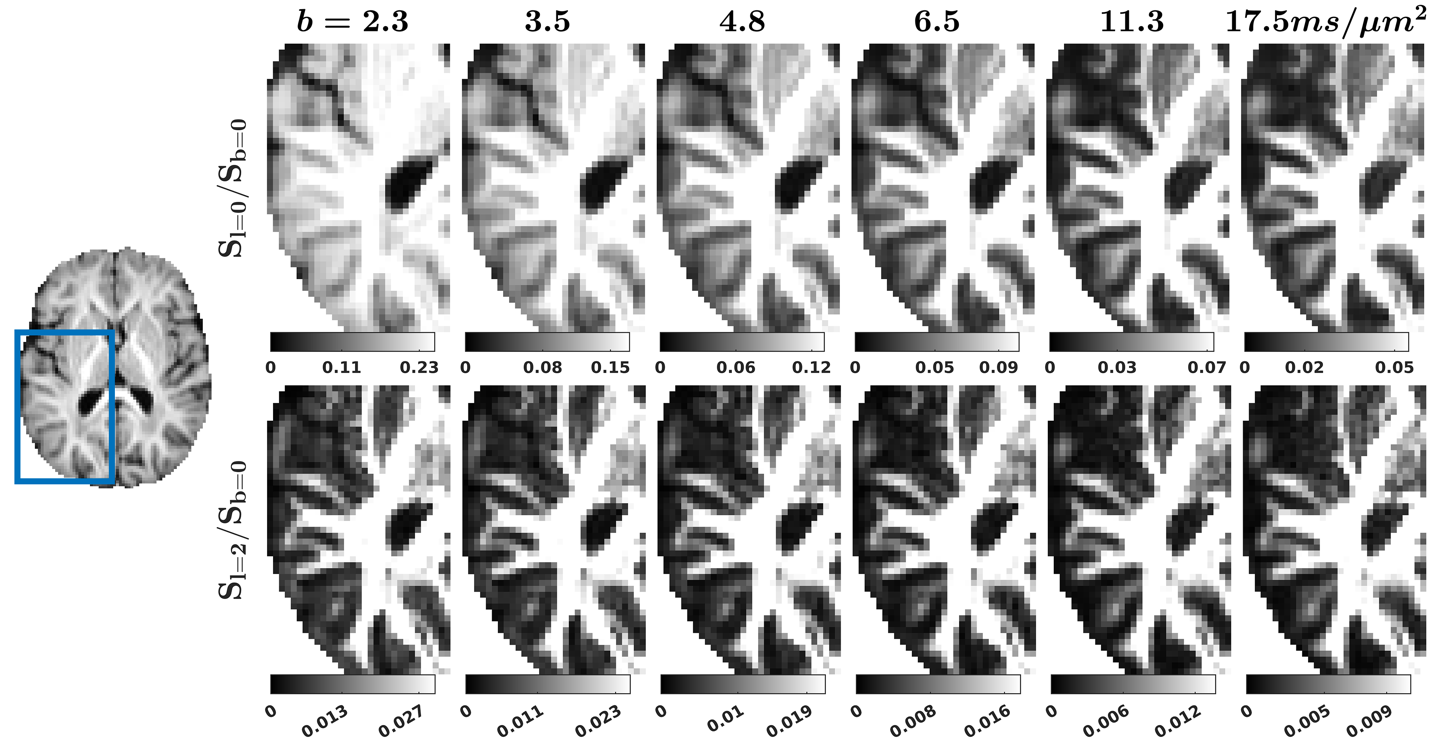

Figure S8: Zoom-in views of the zeroth and second order rotationally invariant dMRI signals at t of 30 ms and *b*-values from 2.3 ms/µm^2^ to 17.5 ms/µm^2^ on one subject from the high SNR Connectome 2.0 cohort on the same slice shown in Figure 6.

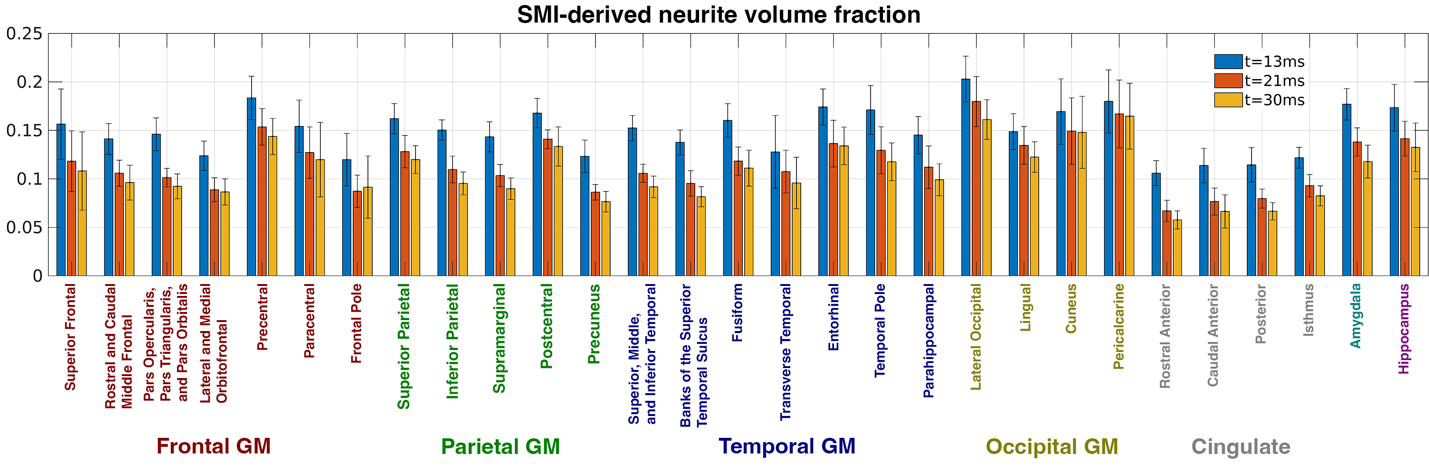

Figure S9: Neurite volume fraction across various cortical ROIs and across all subjects without considering the exchange effect on the Connectome 2.0 dataset. Voxel-wise estimation was performed using the Standard Model Imaging (SMI) toolbox (Coelho et al., 2022) on dMRI data acquired at three diffusion times, respectively.

**Reference**

Coelho, S., Baete, S.H., Lemberskiy, G., Ades-Aron, B., Barrol, G., Veraart, J., Novikov, D.S., Fieremans, E., 2022. Reproducibility of the Standard Model of diffusion in white matter on clinical MRI systems. NeuroImage 257, 119290. <https://doi.org/10.1016/j.neuroimage.2022.119290>
